# Epiblast Lumenogenesis is not a mammalian-specific trait

**DOI:** 10.1101/2025.08.06.669009

**Authors:** Antonia Weberling, Natalia A. Shylo, Hannah Wilson, Melainia McClain, Richard Kupronis, Alex Muensch, Suzannah A. Williams, Florian Hollfelder, Paul A. Trainor

## Abstract

Epiblast lumenogenesis, which leads to amniotic cavity formation, is a hallmark of mammalian embryogenesis, and required for crucial developmental processes such as anterior-posterior patterning and gastrulation. Based on avian model-organisms, the epiblast in reptiles is thought to form a monolayered flat disc that undergoes anterior-posterior patterning and gastrulation. Here, we report that the squamate, veiled chameleon (*Chameleo calyptratus*), exhibits epiblast lumenogenesis and amniogenesis prior to anterior-posterior patterning. Using SEM, immunofluorescence, and histology techniques, we demonstrate that chameleon epiblast lumenogenesis occurs via a purse-string-like mechanism involving the formation of supracellular actin cables in concentric rings around the epiblast followed by constriction that closes the epiblast lumen. Through expression analyses of *Nodal1*, *Nodal2*, *Cerberus*, *Lefty*, *Brachyury, Wnt3A*, and *Bmp2*, and immunostaining for Brachyury, we uncovered a *Wnt3a-* and Brachyury-positive ring at the edge of the epiblast concomitant with lumenogenesis, and preceding anterior-posterior patterning and gastrulation. Furthermore, we report anterior-posterior patterning in veiled chameleons occurs independently of *Cerberus* and *Lefty*. These processes that mediate epiblast lumenogenesis in chameleons, result in a morphology remarkably similar to human embryos, despite 300 million years of evolutionary separation. Taken together, we show that pre-gastrulation epiblast lumenogenesis is not mammalian-specific, but also occurs in some non-avian reptiles.

## Introduction

Epiblast lumen formation is a hallmark of mammalian pre-gastrulation morphogenesis and essential for amnion formation, anterior-posterior development, and gastrulation.^1^ In human embryos, the epiblast lumen forms upon implantation through charge repulsion and fluid pumping in concert with cell shape changes and dynamic specific gene activity.^2,3^ The initially naïve epiblast consists of unpolarised pluripotent embryonic stem cells that subsequently establish apical-basal polarity and form a monolayered epithelium surrounding a central lumen. Simultaneously, these stem cells transition from naïve pluripotency to a primed state.^4,5^ The proximal side of the epiblast then becomes squamous and differentiates into the amnion.^6,7^ Epiblast lumenogenesis has been reported to be critical for anterior-posterior patterning and initiation of gastrulation in mice with Bmp4 being secreted by the extraembryonic ectoderm into the proamniotic cavity to induce Wnt3 and Nodal expression in the epiblast.^1^ This in turn defines the distal visceral endoderm (DVE), which expresses Nodal antagonists Cerberus1, Lefty1, Goosecoid (Gsc) and Hex.^8^ The anterior-posterior axis is established through migration of the DVE to the epiblast-extraembryonic ectoderm border.^8^ In human embryos, CERBERUS1 and LEFTY1 were recently shown to establish an anterior hypoblast signalling centre.^9^ However, the full signalling cascade remains to be uncovered as the extraembryonic trophoblast layer is not in contact with the amniotic cavity.

The chicken embryo has served as a primary organism for modelling (pre-)gastrulation during human embryogenesis and is also thought to represent avian and non-avian development more generally.^10^ During chicken embryogenesis, the epiblast and hypoblast form a 2-dimensional flat bilaminar disk through thinning of a multilayered blastoderm.^11–13^ The anterior-posterior axis is established during this thinning process, through Nodal signalling and the expression of its antagonists.^14,15^ However, the chicken embryo lacks lumenogenesis. The veiled chameleon (*Chameleo calyptratus*), a scaled, non-avian reptile (squamate) species is an emerging model for studying reptile gastrulation and the evolution of developmental mechanisms.^16–18^ Unlike other squamates, chameleon embryos are pre-gastrula stage at the time of oviposition, ^16,18,19^ and laid in clutches of up to 90 roughly time-matched eggs year-round^20^. The veiled chameleon genome was recently annotated^21^ and karyotyped^22^ making it a genetically tractable model organism. While its post-oviposition development gains increasing attention, chameleon pre-oviposition development was last studied in the 1930s.^23,24^

Here, we describe chameleon pre-oviposition development and reveal that unlike other reptiles, chameleons form an epiblast lumen. Thereby the chameleon embryo morphology becomes remarkably similar to human embryos^7^ with a dorsal trophoblast-like layer, a ventral hypoblast layer and an epiblast surrounding a central lumen with the dorsal portion of the epiblast exhibiting squamous, amnion-like morphologies (Figure 1A). We uncovered circular Brachyury expression and an epithelial-to-mesenchymal-like transition coinciding with lumenogenesis, prior to anterior-posterior patterning, raising the question of when gastrulation initiates. Lumenogenesis in veiled chameleons appears to be mediated via a purse-string mechanism with supracellular actin cable constriction. Lastly, we also describe the evolutionary divergence of anterior-posterior patterning between chameleons, and mammals and chicken. Collectively our work reveals that lumenogenesis and pre-gastrulation amnion formation are not mammal specific and may have evolved multiple times.

**Figure 1:**
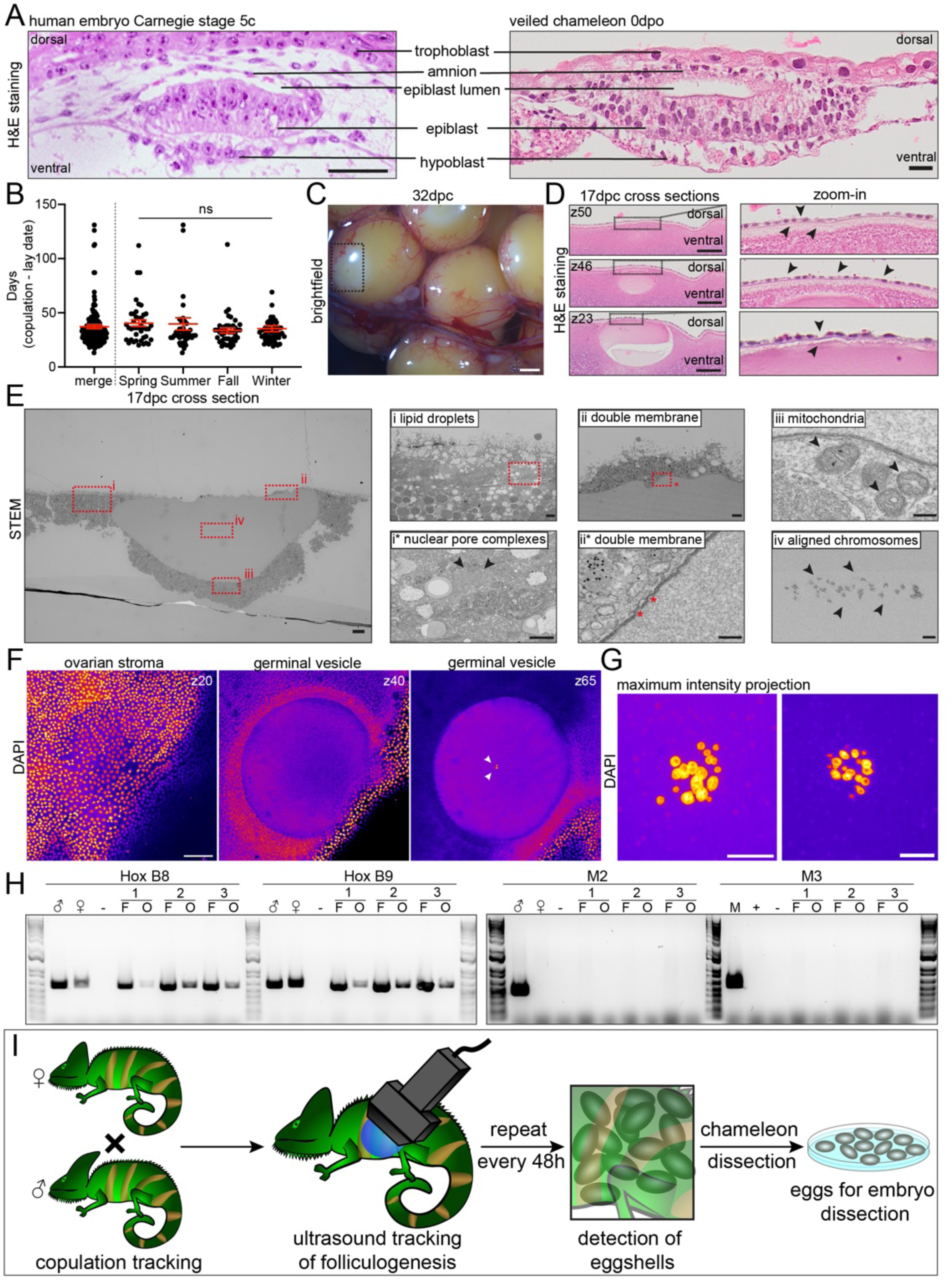
Germinal Vesicle morphology in veiled chameleons. **A.** left H&E stainings of cross section of Carnegie stage 5c human embryo^7^ (left) and 0dpo veiled chameleon embryo (right). dorsal-ventral axis of the embryos is annotated. The position of trophoblast, amnion, epiblast lumen, epiblast, and hypoblast are indicated. Scale bar human=50μm. chameleon=20μm. **B.** Timing of pre-oviposition development in the veiled chameleon. Scatter plot of overall time between mating and oviposition (left) and separated by season (right). Statistical analysis median±SEM: combined: 32±1.89 days, spring: 36±3.03, summer: 29.5±5.73, fall: 32±2.35, winter: 32±2.80. One-way Anova: p-value=0.5063. **C.** brightfield image of mature vitellogenic follicles in ovary at 32dpc. box: germinal disc with germinal vesicle. Scale bar: 2mm. **D.** H&E staining of cross sections of a germinal disc. The germinal vesicle appears as empty sphere. Zoom in on the monolayered ovarian stroma. Scale bars: 200μm. **E.** TEM of germinal vesicle. Red squares indicate zoom ins. Scale bar: 20μm. (i) the yolk around the germinal vesicle is rich in lipid droplets, (i*) nuclear pore complexes and golgi apparatus, (ii) germinal vesicle surrounded by double membrane, (ii*) double membrane with pores, (iii) mitochondria found in direct proximity to the germinal vesicle, (iv) aligned chromosomes in middle of germinal vesicles. Scale bars: i/ii/iv: 2um, i*: 1um, ii*/iii: 200nm. **F.** 3D confocal imaging of DAPI-stained germinal vesicle. DNA is visible in the middle of the germinal vesicle (z65) Scale bar: 100μm. **G.** Spinning Disc imaging of DNA in centre of germinal vesicle. The DNA appears separated into single chromosomes. Scale bars 10μm. **H.** Genotyping of DNA extracted from follicles (F) and germinal vesicles (O) of 3 individual clutches (1-3). Control genes: HoxB8, HoxB9, male loci M2, M3. As controls male DNA (^♂^), female DNA (^♀^) and a water control (-) were used. **I.** Schematic of experimental set-up. Following copulation, chameleons are subjected to ultrasound every 48h. Upon detection of specific eggshell maturity, chameleons are dissected. Then the embryos are dissected and analysed.

## Results

### Sperm storage leads to variable time between copulation and oviposition

To study chameleon pre-oviposition morphogenesis, it was necessary to first define the timing of key reproduction and early embryo developmental events. Staging conventions are commonly based on time passed since copulation which correlates with fertilisation.^1,13,25^ Chameleons lay 3-4 clutches per year,^20^ and by monitoring the copulation and oviposition dates we observed an average lay date of 37 days post copulation (dpc) (Figure 1B). We decided to isolate eggs between 17-37 dpc but could only observe large vitellogenic follicles in the ovaries (Figure 1C, S1A). Each follicle contained a germinal disc with the germinal vesicle in the middle (Figure 1C, box, S1A, boxes asterisks). Histological sections of the germinal disc shows that the germinal vesicle (∼300*μ*m diameter) (Figure 1D) is overlaid by the ovarian stroma (Figure 1D, arrows). We then carried out TEM of the germinal vesicle (Figure 1E). The yolk is very rich in lipid droplets (Figure 1Ei) and we could observe nuclear pore - and Golgi complexes (Figure 1Ei*). The nuclear envelope exhibits nuclear pores (Figure 1Eii/ii*) and has mitochondria in the immediate proximity (Figure 1Eiii). Interestingly, we observed distinct structures in the middle of the germinal vesicle reminiscent of aligned chromosomes (Figure 1Eiv). More than 20 structures could be counted, which was suggestive of a diploid set of chromosomes, since the chameleon has 24 chromosomes.^21,22^ To confirm this observation, we DAPI stained the germinal discs and found that this central structure retained the DAPI label (Figure 1F z65, S1B z66, arrows) and was composed of over 12 DAPI-positive foci (Figure 1G, S1C), again suggesting diploidy. These follicles were collected at various timepoints post mating suggesting that fertilisation timing may not correlate with copulation. Squamates are known for short- and long-term sperm storage^26,27^, but this had not been previously described for veiled chameleons. To confirm that these follicles were indeed oocytes we dissected the germinal vesicles of three clutches at 30-37dpc and extracted DNA of the pooled germinal vesicles (O) and immature follicles (F). Genotyping for two control genes (HoxB8/9)^28^ confirmed successful DNA extraction (Figure 1H left). Negative genotyping for two male-specific loci (M2/3)^21^ revealed that the follicles are indeed oocytes (Figure 1H right) confirming that chameleons store sperm. Thus, the day of copulation cannot be used to calculate embryonic age. However, eggshell formation initiates with embryogenesis, and density of the eggshells compared to surrounding inner organs can be detected via ultrasound and used to predict pre-oviposition stage^29^ (Figure S1D). The eggshell is initially only faintly visible (Figure S2C) but matures into a clearly distinguishable ridge increasing in width and thickness (Figure S2D-F). We therefore used ultrasound imaging to monitor embryo progression enabling our investigation of pre-oviposition embryogenesis (Figure 1I).

### Cleavage divisions give rise to a multilayered embryonic shield

The initial cleavage furrows form following fertilisation in eggs covered with very thin eggshells (Figure S3A/B). The first furrow spans the length of the embryonic plate which can be discriminated from the germinal disc by different surface structures (S2B blue). The next cleavages initiate through furrows emanating from the embryo centre (S3Bi*) and holes at the border of the embryonic plate (S3Bii**/***). This pattern results a radial initial cleavage pattern within the embryonic plate (Figure 2A,Ai-ii*). The embryonic plate border can be clearly observed through SEM (Figure 2Aiii/iii*) and F-actin and WGA labelling (Figure 2B top). Staining with DAPI, revealed individual DAPI-positive foci in the individual blastomeres but no enlarged spheres as in the germinal vesicle. (Figure 2B, bottom, arrows). TEM of single blastomeres revealed the nucleus localises to the dense dorsal region (Figure S3C, i). The staining inside the nuclear envelope, exhibits characteristic density patterns suggestive of multiple nucleoli Figure 2C, S3Ci*). The double-layered envelope contains nuclear pores (Figure S3Ci88) and is surrounded by mitochondria and Golgi (Figure S3Cii-ii*). Interestingly, we observed a high concentration of filaments oriented in different directions in association with cleavage furrow formation (Figure 2Cii, S3ii**,iii-iv).

**Figure 2:**
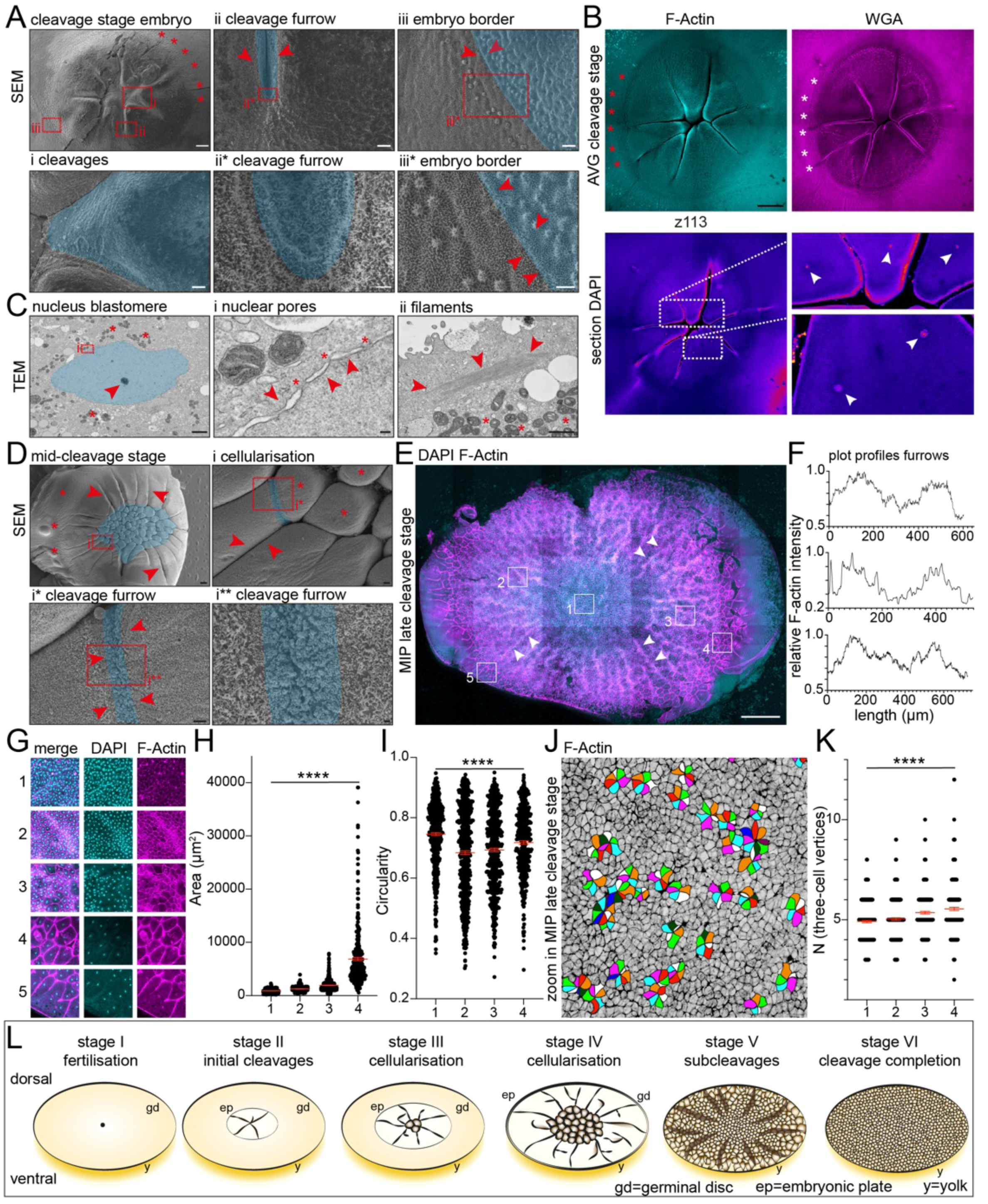
Initiation and progression of Cleavage divisions. **A.** SEM imaging of early cleavage stage embryo. asterisks indicate embryo plate border. (i) cleavages, blue highlight initial blastomere. (ii/ii*) cleavage furrow highlighted in blue. (iii/iii*) embryo border, embryonic plate in blue, arrows indicate border. scale bars: cleavage stage embryo=200μm, i/ii/iii = 40μm, ii*=5um, iii*=20μm. **B**. Confocal imaging of early cleavage stage embryo. top: average intensity projection, left F-actin, right Wheat-Germ-Agglutinin. Embryo border highlighted with asterisks. bottom: DAPI staining of z-slice z113. Individual nuclei of nascent blastomeres indicated with arrows. scale bar:400μm. **C.** TEM imaging of initial cleavage blastomere. Left nucleus blastomere. Nucleus highlighted blue, arrow indicates nucleolus. Asterisks indicate mitochondria. (i) Nuclear pores. Double membrane indicated by arrows, pores by asterisks. (ii) Filaments. Filaments highlighted with arrows, mitochondria by asterisks. scale bars: nucleus blastomere=2um, i=100nm, ii=1um. **D.** SEM imaging of mid-cleavage stage embryo. Cellularised area in blue. arrows indicate large cleavage furrows, asterisks border of the embryonic plate. (i) Cellularisation. Asterisks indicate cellularised blastomeres, arrows mature cleavage furrows, immature cleavage furrow in blue. (i*/**) Cleavage furrow. Immature cleavage furrow in blue, highlighted with further arrows. Scale bars mid-cleavage stage 200μm, i=20μm, i*=10μm, i**=2um **E.** Confocal Imaging of late cleavage stage embryo. maximum intensity projection, DAPI (cyan), F-Actin (magenta). arrows indicate large furrows. boxes 1-5 illustrate areas of the embryo. scale bar:1mm. **F.** Plot profiles of F-actin intensity across cleavage furrows. Each plot profile was drawn in 100μm thickness across 2 cleavage furrows illustrating the difference in actin intensity within and between the furrows. **G.** zoom-ins of (E) DAPI (cyan) & F-Actin (magenta) staining of different areas of late cleavage stage embryo. **H.** Analysis surface area of embryonic cells of areas 1-4. Scatter plot of 3 embryos per area. Scatter plot plus mean±SEM. Statistical analysis: One-way Anova p-value<0.0001, unpaired t test areas 1-2, 2-3, 3-4, each p-value <0.0001. **I.** Quantitative Analysis of embryo cell circularity across areas 1-4. 3 embryos were analysed. Scatter plot plus mean±SEM. Statistical analysis: One-way Anova p-value<0.0001, unpaired t test areas 1-2 p-value <0.0001, areas 2-3 p-value=0.3012, areas 3-4 p-value=0.0048. **J.** Rosettes within the centre of late cleavage stage embryos. **K.** quantitative analysis of three-cell vertices across areas 1-4. 3 embryos were analysed. Scatter plot plus mean±SEM. Statistical analysis: One-way Anova p-value<0.0001, unpaired t-test areas 1-2 p-value=0.1323, areas 2-3 p-value=0.0002, areas 3-4 p-value=0.0606. **L.** schematic of Embryogenesis across cleavages. Stage I fertilisation. Stage II initial cleavages. The initial cleavage furrows form inside the embryonic plate (ep) that is located in the middle of the germinal disc (gd). Stage III cellularisation. The initial blastomeres form in the middle of the embryonic plate, which extends further over the germinal disc. Stage IV: cellularisation. the embryonic plate covers the entire germinal disc, the centre of the embryo is filled with large blastomeres, large cleavage furrows emanate from the blastomeres to the edges of the embryonic plate. Stage V: subcleavages. The embryonic plate is filled with blastomeres which are small in the centre and larger towards the outside. Large furrows go through the entire embryo. stage VI: cleavage completion. The entire embryonic plate is filled with small blastomeres. Gd=germinal disc, ep=embryonic plate, y=yolk.

Subsequently, the first blastomeres form while the cleavage furrows span the entire embryonic plate, which by now extends over the entire germinal disc (Figure 2D/S3D). The cleavage divisions continue until the entire embryonic plate is composed of blastomeres (Figure 2E). Interestingly, large radial furrows extend from the middle towards the edge of the embryonic plate even though the entire plate is already composed of blastomeres, similar to *Anolis sagrei*^30^ (Figure 2E arrows). These furrows exhibit high actin intensity indicative of tension^31^ (Figure 2F). During this period, the eggshell becomes more pronounced (Figure S3E). Within the late cleavage stage embryo, five different areas can be distinguished. In the middle (Figure 2E/G 1) the blastomeres appear dense. In the adjacent area (2), the cells are less crowded but still homogenous. Radiating further laterally, both large and small blastomeres are found in area 3, which become even larger more laterally in area 4. Area 5, the most lateral domain, is in direct contact with the edge of the embryonic plate and contains open cleavage furrows and multinucleated cells (Figure 2G5, arrows). We validated these qualitative observations through quantitative analysis showing exponential increase of cell area and perimeter while revealing high variability in cell circularity. Central blastomeres exhibit high circularity, which decreases in blastomeres located more laterally in area 2, before increasing again in areas 3 and 4 (Figure 2H/I, S4A-C). During our analysis, we observed cells arranged in multicellular rosettes (Figure 2J), a sign of increased tissue rigidity.^32^ Epithelia exhibit hexagonal packaging,^33^ and we observed an overall increase in three-cell vertices from area 1-5. This may be accounted for by the increase in size and perimeter (Figure 2K, S4D). Taken together, we define 6 embryonic stages during the initial stages of chameleon embryogenesis leading from fertilisation (stage I) to the completion of cleavage divisions with an embryo formed of small non-adherent blastomeres (stage VI) (Figure 2L, S5).

### Epiblast lumenogenesis occurs via tissue folding

Following the completion of cleavage divisions, the blastomeres become adhesive and form a coherent sheet^30^ (Figure 3A, S6A). Tightly packed cells build the dorsal side of the embryo (Figure 3Ai-iii, arrows), while the ventral side exhibits larger, loosely packed, rounded cells (Figure 3A/S6A asterisks, 3B/C, S6B/C). This ventral population thickens in the middle of the embryo (Figure 3A, S6A) and is more apparent through immunofluorescent imaging which illustrates the crowding of the nuclei towards the middle of the embryonic shield (Figure 3D). The dorsal cells exhibit comparable surface areas throughout the embryo, thus, the higher density of nuclei in the middle must be associated with the ventral cell layers (Figure 3B, S6D). These cell populations give rise to the epiblast (dorsal) and the hypoblast (ventral). Subsequently, the embryo which was initially a flat tissue, hollows and forms a dome-like structure (Figure 3E) similar to the brown anole.^30^ Two shoulders of tissue on each edge of the epiblast (Figure 3Ei, arrows), appear thicker than the remaining epiblast (Figure 3Eii). Subsequently, the side shoulders fold dorsally over the dome-shaped epiblast (Figure 3F arrows). This tissue folding coincides with apical constriction in the hinge points cells (Figure 3Fi). The cells at the tips of the annealing folds orient towards the tissue surface (Figure 3Fii, asterisks), and while the folds anneal further but are not yet closed (Figure 3G arrows), the epiblast curvature inverts from concave to convex (Figure 3E-G). A squamous cell layer is then observed dorsally (Figure 3G/i-ii highlight) while the remaining fold cells appear disorganised. The embryo is embedded in a multilayered tissue originally located at the outer area of the embryonic shield (Figure 3Gii). Simultaneously, the hypoblast rearranges from a thick, spongy tissue into a thin cell layer surrounding the convex epiblast (Figure 3Eii, F, Giii asterisks).

**Figure 3:**
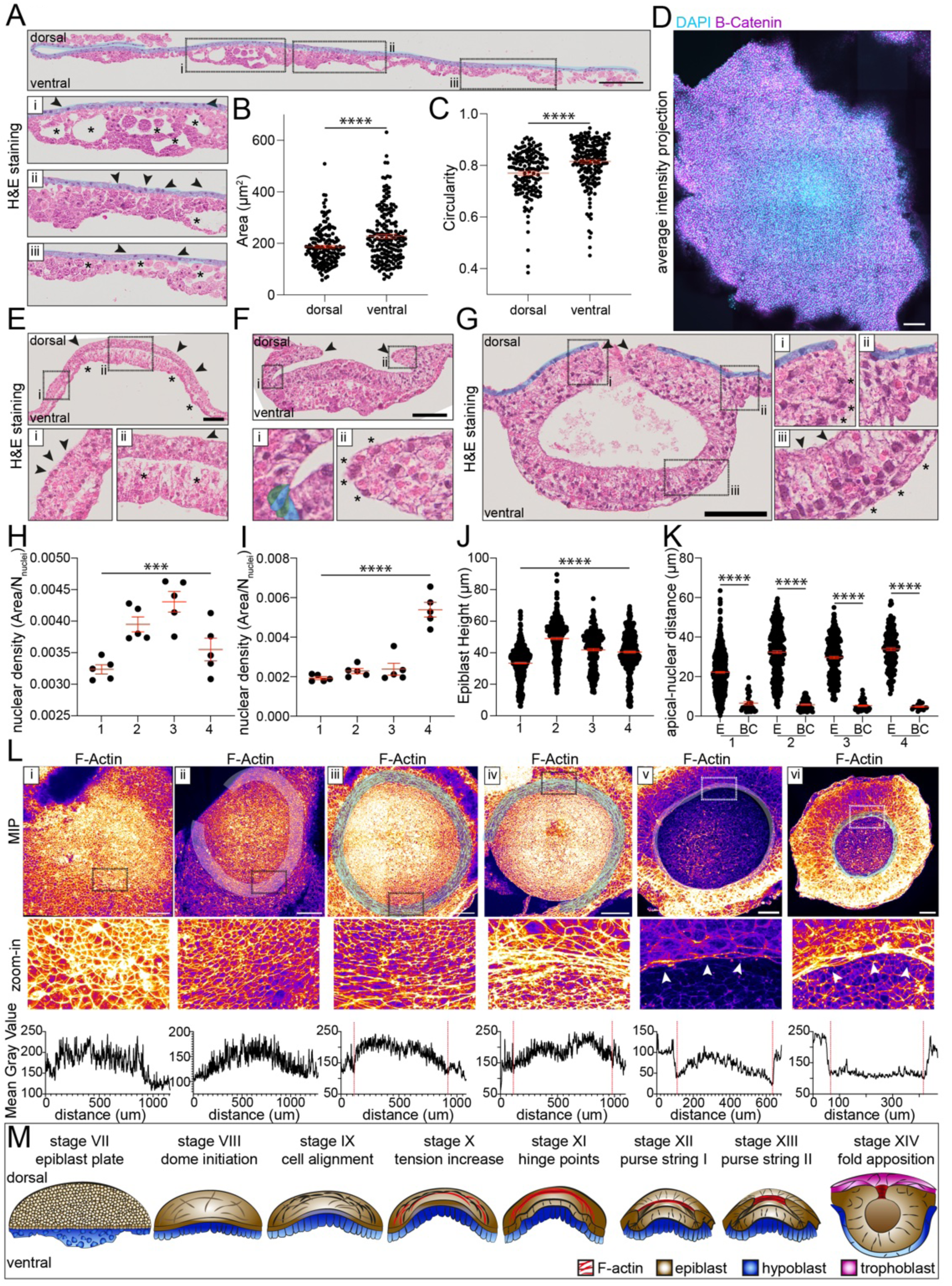
The Epiblast Lumen forms via a purse-string mechanism. **A.** H&E staining of post-cleavage stage embryo. The embryo is composed of a coherent dorsal tissue composed of squamous or cuboidal cells, the prospective epiblast (blue) and a spongy ventral tissue, the prospective hypoblast. (i-iii) zoom in of different embryo regions epiblast in blue, and indicated by arrows, the spongy hypoblast indicated with asterisks. scale bar=200μm. **B-C.** Quantitative analysis of dorsal versus ventral tissues for Area (B) and Circularity (C), 3 embryos were analysed. Scatter plot plus mean±SEM. Statistical analysis unpaired t-test p-values area<0.0001, circularity<0.0001. **D.** Confocal Imaging of post-cleavage stage embryo. Average intensity projection of DAPI (cyan) and β-Catenin (magenta). scale bar=200μm. **E-G.** H&E stainings of embryos as successive lumenogenesis stages. squares indicate region of zoom-in. (E/ii) arrows indicate epiblast, asterisks hypoblast. (Ei) arrows indicate hinge-point. (F) amnion folds indicated by arrows. (Fi) apically constricted cells in hinge-point. (Fii) asterisks indicate border cells. (G) amnion fold apposition (arrows) trophoblast-like layer indicated in blue. (Gi) asterisks apposing fold cells. (Giii) asterisks indicate hypoblast, arrows epiblast. All scale bars 100μm. **H/I.** quantitative analysis of nuclear density (tissue area/number of nuclei within tissue) of epiblast (H) and Hypoblast (I) during successive stages of lumenogenesis (1-4). Scatter plot, mean±SEM. Epiblast: Statistical Analysis one-way Anova p=0.0004, unpaired t-tests p-values (1-2)=0.001, (2-3)=0.1156, (3-4)=0.0148. Hypoblast: Statistical Analysis one-way Anova p<0.0001, unpaired t-tests p-values (1-2)=0.0483, (2-3)=0.7716, (3-4)=0.0002. **J.** Epiblast tissue height during successive stages of lumenogenesis (1-4). Scatter plot, mean±SEM. Statistical analysis one-way Anova p<0.0001, unpaired t-tests p-value (1-2)<0.0001, (2-3)<0.0001, (3-4)=0.1527. **K.** apical-nuclear distance in the epiblast (E) versus the tip cells in the lumenogenesis folds (BC) during lumenogenesis. Scatter plot, mean±SEM. Statistical analysis unpaired t-tests of E vs BC all p-values<0.0001. **L.** top row maximum intensity projections of F-actin during lumenogenesis (i-vi) in whole embryos. squares indicate zoom-in regions of the second row. white/blue highlights indicate epiblast border. Columns v/vi arrows indicate supracellular actin cable. Bottom row. Plot profiles of actin mean gray value drawn across the length of the embryo. red lines indicate border between epiblast and fold/outer cells. Fire staining (yellow-white: high signal intensity, purple-black low signal intensity). Scale bars: (i-iv)=200μm, (v-vi)= 100μm **I.** schematic of embryogenesis during lumen formation comprising stages VII-XIV. Epiblast plate (stage VII), dome initiation (stage VIII), cell alignment (stage IX), tension increase (stage X), hinge point initiation (stage XI), the start of the purse string (stage XII), constriction of the purse-string (stage XIII), and fold apposition (stage XIV). F-actin (red), epiblast (brown/beige), hypoblast (blue), and trophoblast-like cells (magenta) are included in the schematic.

Measuring nuclear density in epiblast and hypoblast at successive stages of lumenogenesis reveals an increase in epiblast nuclear density during folding, which diminishes upon fold annealing (Figure 3H). The hypoblast nuclear density remains constant and only increases dramatically upon inversion of epiblast curvature (Figure 3I). Epiblast height is higher in the middle versus the sides and increased slightly during folding but then decreases upon fold annealing (Figure 3J). The apical-nuclear distance of epiblast cells is highly variable, indicative of a pseudostratified epithelium while the tip cells exhibit much shorter distances to the apical surface (Figure 3K).

We next investigated lumenogenesis through whole mount F-actin staining (Figure 3L). Upon the initiation of hollowing, the central epiblast cells were significantly smaller than outer cells and exhibited higher actin intensity and higher circularity (Figure 3Li, S7A-I) This size difference remained throughout lumenogenesis while the circularity overall decreased. Once the epiblast dome matured, the border cells aligned in concentric rings around the epiblast centre (Figure 3Lii). Then, the border cells elongated and exhibited slightly higher actin levels (iii) until supracellular actin rings appeared around the epiblast dome in subsequent stages (Figure 3Liii-iv, red lines plot profiles). The ring-cells stretched so thin, that individual cells couldn’t be distinguished (Figure 3Liv/v). Subsequently, these ring-cells disappeared and instead, the epiblast was overlaid and surrounded by an actin ring in direct contact with the large outer cells (Figure 3Lv, arrows). The epiblast could be distinguished from the outer cells on top by size and different actin levels (Figure 3Lv plot profile, red lines). The outer cells then continuously constrict further over the epiblast exhibiting even higher actin intensity than the underlying epiblast (Figure 3Lvi). Taken together, we define 8 consecutive stages of lumenogenesis stages from the coherent embryo (stage VII) to a the closely annealed, although not-yet closed epiblast lumen (stage XIV) (Figure 3M, S5).

### Epiblast lumen closure is followed by symmetry breaking and gastrulation

The epiblast lumen closes via the purse-string with the closure point remaining visible (Figure 4A/Ai). The epiblast cells are columnar (Figure 4Aii) and overlie the hypoblast, a loosely packed pseudostratified epithelium (Figure 4Aiii). The cell layer in-between the epiblast and the emerging trophoblast appears disorganised (Figure 4Aiv). Upon oviposition, the trophoblast-like tissue transforms into enlarged, squamous cells overlaying highly squamous amnion-like cells (Figure 4B/Bi arrows) with the closed lumen still visible as a morphological tissue thickening (Figure 4B, blue). Gastrulation is visible by spindle-shapes cells that delaminate from the epiblast and migrate between the epiblast and hypoblast (Figure 4Bii). The epiblast itself undergoes posterior thickening through apicobasal elongation of the epithelium (Figure 4Biii/iv asterisks). Subsequent embryo symmetry breaking and the initiation of gastrulation can be followed through SEM (Figure 4Ci-iii).

**Figure 4:**
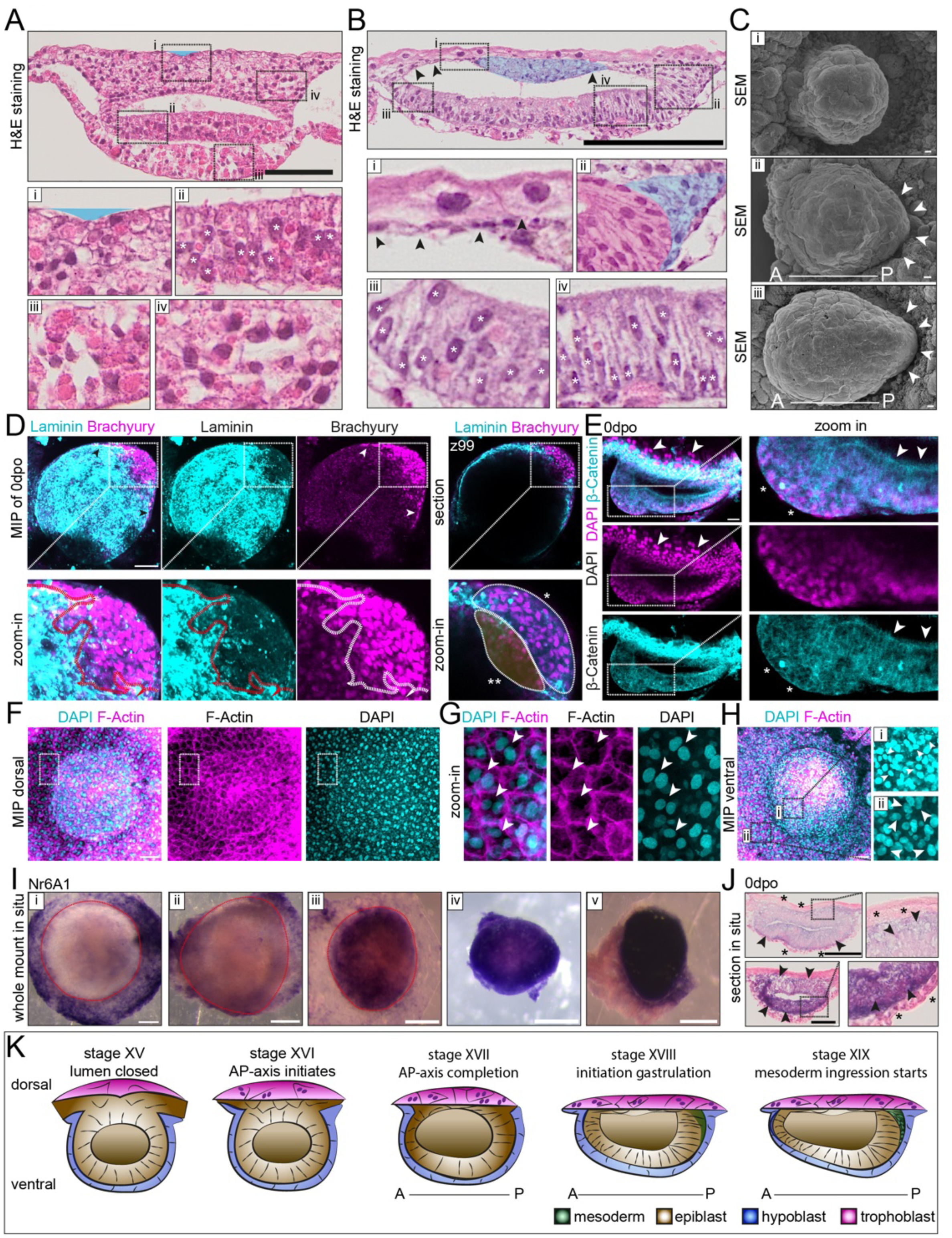
Lumenogenesis is followed by anterior-posterior patterning and gastrulation. **A.** H&E staining of preoviposition embryo following lumen closure. The amniotic fold fusion point visible as dorsal indentation (blue highlight, i). zoom ins: (i) amniotic fold fusion (ii) zoom in epiblast, asterisks indicate nuclei (iii) hypoblast (iv) epiblast-derived dorsal tissue shows no polarity. scale bar=200μm. **B.** H&E staining of 0dpo embryo. amnion fusion remnant highlighted in blue. squamous amnion cells indicated by arrows. zoom-ins: (i) dorsal enlarged trophoblast-like cells. ventral squamous amnion (arrows). (ii) posterior embryo and mesoderm ingression. Embryo highlighted in magenta, ingressing mesoderm in blue. (iii/iv) anterior (iii) and posterior (iv) epiblast with nuclei highlighted through white asterisks. scale bar=200μm. **C.** SEM imaging of symmetry breaking. Imaging from ventral side. Arrows indicate posterior. scale bars: i/iii=20μm, ii=30μm. **D.** 0dpo embryo, confocal imaging of IF staining of Laminin (cyan) and Brachyury (magenta). left 3 columns maximum intensity projection, zoom-ins: dashed line indicates breach of basement membrane. Right column single z-section. Zoom-in: *-marked outline ingressed mesoderm cells, **-marked outline mesoderm cells that have not ingressed through basement membrane yet. scale bar=100μm. **E.** 0dpo embryo, confocal imaging of DAPI (magenta) and β-Catenin (cyan) immunofluorescence staining. Arrows indicated double-nucleated trophoblast-like cells. zoom ins: arrows indicate pseudostratified epithelium in epiblast. asterisks indicate rounded mesoderm cells following ingression. scale bar=50μm. **F.** maximum intensity projection of dorsal side of embryo following lumen closure, DAPI (cyan) and F-actin (magenta). squares indicate zoom ins of (G), scale bar=100μm. **G.** zoom ins of trophoblast-like cell layer (F). arrows indicate double nucleated cells. **H.** maximum intensity projection of ventral side of embryo following lumen closure. DAPI (cyan) and F-actin (magenta) squares indicates zoom ins (i/ii). i. zoom in on hypoblast/epiblast nuclei, arrows indicate single nuclei. ii. zoom-in on trophoblast-like nuclei. Arrows indicate single nuclei. Scale bar=100μm. **I.** whole mount in situ hybridisations of *Nr6a1* from initiation of lumenogenesis to oviposition. Red line indicates epiblast. all scale bars=200μm. **J.** cross section of in situ hybridisation for *Nr6a1* of 0dpo embryo. dorsal top. Arrows indicate epiblast. asterisks indicate extraembryonic tissues. Scale bars: top 200μm, bottom 100μm. **K.** schematic of embryogenesis from lumen formation to the initiation of gastrulation comprising stages XV-XIX. Lumen closed (stage XV), AP=axis initiation (stage XVI), AP-axis completion (stage XVII), initiation of gastrulation (stage XVIII), and the initiation of mesoderm ingression (stage XIX). Mesoderm (green), epiblast (beige/brown), hypoblast (blue), trophoblast-like cells (magenta) shown in schematic.

To investigate initiation of gastrulation in more detail, we stained 0dpo embryos for Laminin and the gastrulation marker Brachyury (Figure 4D). Laminin forms a thin basement membrane on the basal side of the epiblast, while a distinct gap at the site of gastrulation was observed (Figure 4D, zoom ins). Brachyury-positive cells ingress out of the epiblast and spread over the laminin (Figure 4D arrows). Two populations of Brachyury positive cells were observed, one migrating out of the epiblast (Figure 4D z99*) and the second within the epiblast (Figure 4D z99**). Gastrulation involves the delamination of cells from the epiblast epithelium, and we therefore analysed 0dpo embryos for β-Catenin, a cell shape marker and DAPI, and report distinct shapes between the highly columnar epiblast, and the smaller, rounded gastrulating cell population (Figure 4F, 4F zoom). Interestingly, the trophoblast-like cell population appeared double nucleated with enlarged nuclei compared to the epiblast/hypoblast populations (Figure 4F-I arrows).

Syncytium-formation and enlargement of nuclei are hallmarks of trophoblast maturation in mammals^34–36^, which this trophoblast-reminiscent population exhibits as well (Figure 4H/I arrows). Since no trophoblast tissue marker was specific, double nucleation and increased size can serve as morphological markers for this trophoblast-like population. We were also not successful in identifying an epiblast protein marker. Instead, we performed in-situ hybridisation for *Nr6a1*, which has been reported to influence the repression of pluripotency.^37–39^ *Nr6a1* was initially expressed in the cells surrounding the doming epiblast, possibly indicating a loss or decrease in pluripotency in these cells (Figure 4J, left). Upon completion of lumen formation, *Nr6a1* expression was downregulated in trophoblast tissue and instead upregulated in the epiblast (Figure 4J, right). Histological sections confirmed *Nr6a1* expression is restricted to the epiblast tissue with negative hypoblast and trophoblast-like cells (Figure 4K). Taken together, we define 5 stages (XV-XIX) between lumen closure and initiation of gastrulation (Figure S5).

### Circular *Brachyury* and *Wnt3a* expression occur prior to anterior-posterior patterning

After developing a morphology-based staging system for functional studies in veiled chameleon, we investigated the signalling pathway dynamics governing anterior-posterior patterning and the initiation of gastrulation. To this end, we carried out *in-situ* hybridisation to define the spatiotemporal expression patterns of the known key regulators *Cerberus*, *Lefty*, *Nodal1*, *Nodal2*, *Bmp2*, *Wnt3a*, and *Brachyury* from stage VIII-XIX. *Cerberus* was expressed as early as stage VIII in a broad domain in the middle of the immature hypoblast (Figure 5A VIII-XIX) similar to human embryos^9^ and remains expressed throughout lumenogenesis. Upon initiation of gastrulation, *Cerberus* expression divides into 2 domains, one remains in the middle of the embryo (Figure 5A XIX circle), while the other forms a distinct domain overlying the site of gastrulation (Figure 5A XIX arrows). Histological sections confirmed that *Cerberus* is expressed in the hypoblast (Figure 5B, arrows) as is characteristic for this gene.^8^ However, *Cerberus* is typically a marker of the anterior but here it is expressed in the posterior of the embryo. Thus, *Cerberus* may not function as a primary regulator of anterior fate or identity in veiled chameleon. *Lefty*, another anterior marker, was also not detected throughout the entire time course (Figure S8B VIII-XIX). Distinct *Lefty* expression during left-right patterning^17^ confirmed that the gene is active but not expressed during anterior-posterior patterning (Figure S8Bi). Focussing on the posterior markers *Nodal1* and *Nodal2*, we observed *Nodal1* expression following lumen formation in a domain demarcating the future posterior region of the embryo, spreading upon gastrulation along the sides of the embryo (Figure 5C, arrows, S8C). *Nodal2* expression initiates at stage IX and localises to one domain on the side of the epiblast dome (Figure 5D). Following lumen formation, *Nodal2* is expressed dorsally in the middle of the embryo, consistent with radial symmetry of the embryo (Figure 5D XV). Subsequently *Nodal2* is expressed in a posterior domain and remains as a small circular domain at the site of gastrulation (Figure 5D stages XVI/XIX). We then examined *Bmp2* expression, which is found in the trophoblast-like tissue from stage VIII until completion of lumenogenesis (Figure 5E, S8D). The posterior marker *Wnt3a* and gastrulation marker *Brachyury* exhibit a striking pattern. *Wnt3a* expression initiates at stage IX in two faint, opposing domains at the edges of the doming epiblast, a pattern replicated by *Brachyury* at stage XI (Figure 5F IX, XI arrows). Upon lumenogenesis, both genes form a dorsal ring over the underlying epiblast (Figure 5F/G XIII) that give rise to dorsal plate of gene expression (Figure S8E XV). Following lumen closure, *Wnt3a* and *Brachyur*y are expressed in two opposing domains (Figure 5F/G XVI). After downregulation in one domain, *Wnt3*a and *Brachyury* localise to the posterior of the embryo and site of gastrulation initiation (Figure 5F/G XVIII-XIX). Cross sections show that *Brachyury* is expressed in the epiblast-derived folds that mediate lumenogenesis (Figure 5H, arrows). Taken together, we observed novel gene expression dynamics of key anterior-posterior axis and gastrulation regulators that deviate from classic mammalian and avian expression patterns.

**Figure 5:**
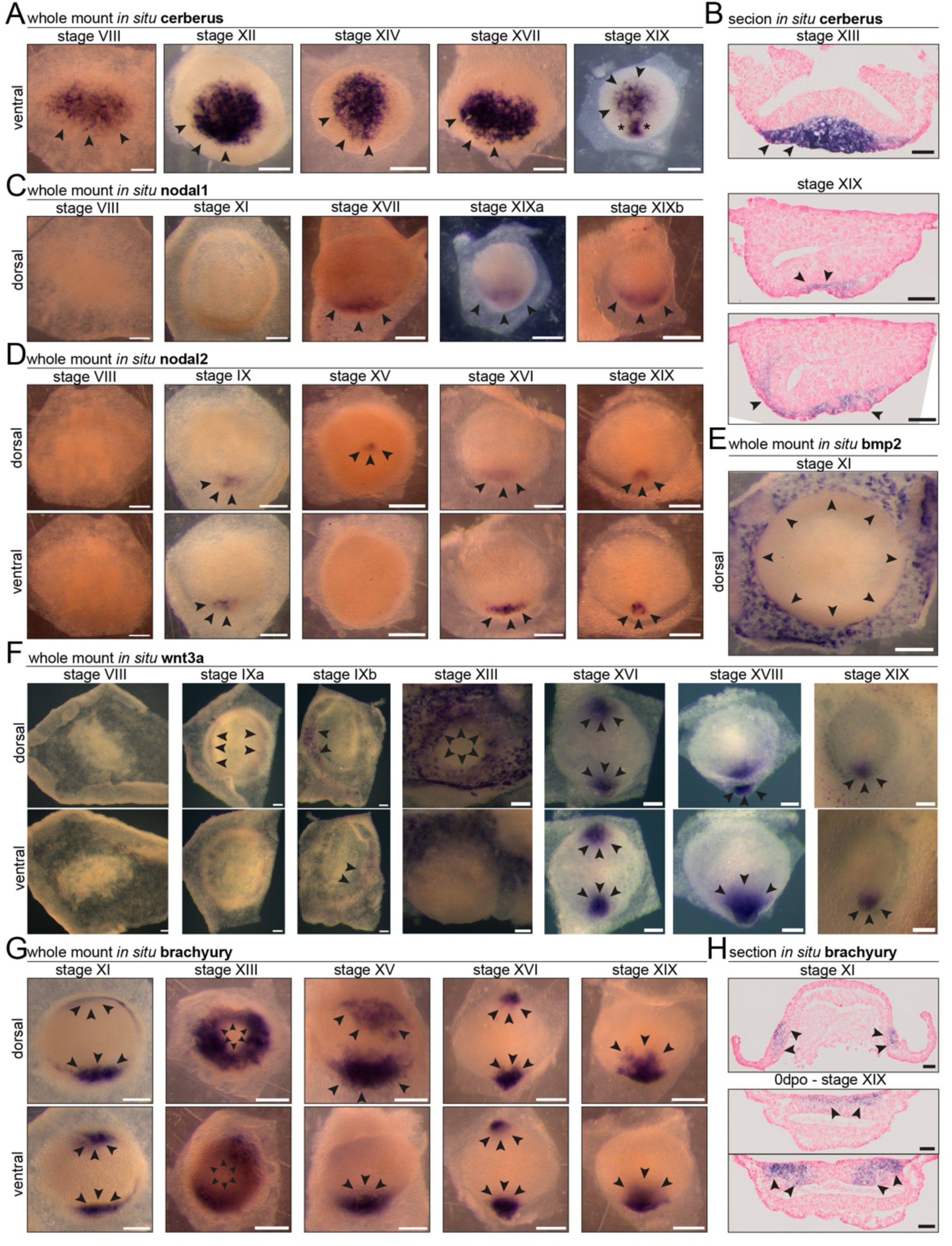
Expression Patterns of anterior-posterior marker genes and gastrulation marker *Brachyury*. **A-H.** gene expression analysis over time (stages VIII-XIX) through in situ hybridisations of **(A/B)** *Cerberus*, **(C)** *Nodal1*, **(D)** *Nodal2*, **(E**) *Bmp2*, **(F)** *Wnt3a*, **(G/H)** *Brachyury*. Arrows indicate sites of expression. asterisks (A, stage XIX) indicate site of gastrulation. Dorsal/ventral views indicated in figure. (A/C-G) whole mount in situs. (B/H) cross sections. Scale bars whole mount in-situs=200μm, scale bars sections= 100μm.

### Circular Brachyury protein expression precedes basement membrane maturation

Our findings raise the question of when gastrulation is initiated in the veiled chameleon. Peter^23^ described mesoderm formation prior to lumenogenesis, however, it is commonly accepted that chameleon embryos are at pre-gastrula or very early gastrula stages upon oviposition.^16,18^ One hallmark of gastrulation is localised basement membrane breakdown enabling ingression of the nascent mesoderm.^40–42^ Following Laminin localisation over time, we initially observed a ring of Laminin strands oriented radially to the centre of the hollowing epiblast (Figure 6A VIII). Laminin then forms thick filaments at the edge of the epiblast dome extending into the trophoblast-like population but not within the epiblast (Figure 6A XI). Upon purse-string initiation, Laminin levels increase in the trophoblast-like population with low levels found in the epiblast (Figure 6A XII). Following lumen closure, Laminin forms a thin basement membrane around the entire, radially symmetric epiblast (Figure 6B XV), which is breached at the site of gastrulation (Figure 6B XIX). Thus, basement membrane maturation follows lumenogenesis and basement membrane breakdown occurs around oviposition. Our *in-situ* hybridisation analyses revealed gene expression patterns but not functional proteins localisation. Therefore, *Brachyury* expression may occur as early as stage VIII, but functional protein may only be present following anterior-posterior patterning. Analysing Brachyury protein expression over time, we observed low levels of Brachyury in scattered cells at the epiblast border upon initiation of epiblast hollowing (Figure 6C VIII). Then, Brachyury activity spreads along half of the epiblast (Figure S9A) developing into a ring with high levels on two opposing sides of the embryos (Figure 6C XI-XII). The Brachyury positive cell population changes morphology from being hexagonal to highly stretched along the amnion folds (Figure 6C zoom-in). Upon lumen closure, Brachyury protein-expressing cells cover the entire dorsal side of the embryo and extend under the trophectoderm (Figure 6D/S9A/B XI-XVIII). We also observed small domains of high Brachyury expression located at the edges of the epiblast. At the onset of gastrulation Brachyury expression was detected only in the posterior of the embryo (Figure 6D XVIII-XIX). Taken together, a circular domain of Brachyury expression prior to basement membrane maturation and anterior posterior patterning, coincides with tissue folding during lumenogenesis.

**Figure 6:**
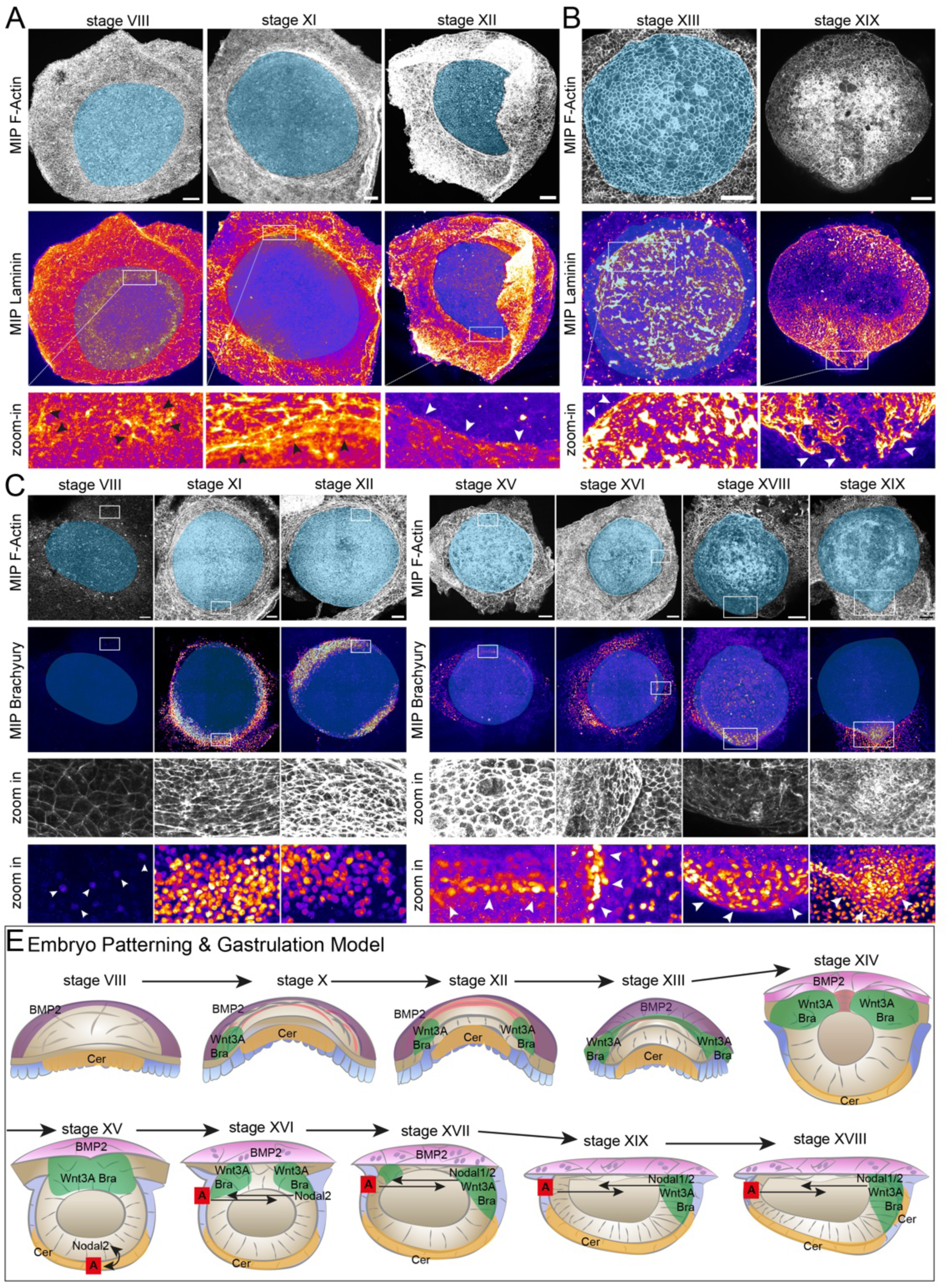
Circular Brachyury protein expression prior to basement membrane maturation A/B. Maximum intensity projection of (A) dorsal view of embryos stages VII-XII. (B) ventral view of embryos stages XIII-XIV. Top row F-actin. bottom row Laminin fire staining illustrating signal intensity (yellow-white: high signal intensity, purple-black low signal intensity). blue highlight indicates epiblast. squares highlight zoom-ins in bottom row. Arrows indicate areas of high laminin intensity (A), or the site of basement membrane breach (B). scale bar embryo1=200μm, all other scale bars 100μm. **C/D.** Maximum intensity projection of (A) dorsal view of embryos stages VIII-XII (B) ventral view of embryos stages XV-XIX. Top row F-actin, Row 2 Brachyury. Blue highlights indicate the epiblast. squares indicate zoom in regions of the bottom 2 rows. Arrows highlight Brachyury positive cells. All scale bars=100μm. **E.** final model of lumenogenesis, anterior-posterior patterning and gastrulation.

## Discussion

Here, we describe veiled chameleon pre-oviposition morphogenesis from fertilisation to gastrulation and the dynamic activity of key regulatory genes governing anterior-posterior patterning and gastrulation. We report several conserved features and vast divergence in epiblast morphology, anterior-posterior patterning and gastrulation.

The initial stages of veiled chameleon embryogenesis are highly conserved between avian and non-avian reptiles.^12,30,43,44^ The apparent presence of nucleoli in the initial blastomeres may suggest active transcription^45^ raising the question of when zygotic genome activation occurs, which happens in chicken at comparable stages (EGK stages I-IV).^46^ During cleavage divisions, the embryo is not cohesive. However, a high number of multicellular rosettes in the embryo centre at late cleavage stages suggest epithelisation.^32^ Once the embryo has become coherent (stage VII), epiblast and hypoblast can be distinguished morphologically, and it will be interesting to investigate how compaction and lineage segregation is mediated during embryogenesis. From stage VII, the chameleon embryo diverges from chicken^11–13^ and other squamates such as brown anoles^30^, asp vipers^44^, and common lizards.^43^

The epiblast undergoes lumenogenesis via a tissue folding and purse-string mechanism to give rise to a structure morphologically very similar to a human embryo.^1,6^ This presents a novel mode of amniotic cavity formation as mammalian epiblast lumens are formed through charge repulsion and hollowing.^2,47^ Following lumenogenesis, the dorsal-most cell layer originating from outer cells of the embryonic plate exhibits enlarged nuclei and binucleation, a hallmark for trophoblast maturation.^34–36^ Thus, the epiblast of early chameleon embryos may have the capacity to give rise to trophoblast-like tissue. While these cells do not form a placenta, they may be a conserved progenitor population.

Human and chameleon embryo morphology appears conserved following lumenogenesis (Figure 1A), however, we report stark divergences regarding signalling dynamics and embryo patterning. Putting all our analysis together (Figure 6E), we propose an alternative anterior gene antagonising *Nodal1/2* and *Wnt3a* to establish the anterior posterior axis in absence of anterior cerberus and lack of lefty expression. Gastrulation has been suggested to occur prior to lumenogenesis^23^ or following lumenogenesis^16,18,19^. While we report Brachyury protein expression at earlier timepoints, we could not observe anterior-posterior patterning which is required for gastrulation prior to lumenogenesis.

These signalling pathway dynamics underlying anterior-posterior patterning and commencement of gastrulation are distinct from mammals suggesting divergent evolution of the anterior-posterior patterning gene regulatory network. To our knowledge, we are the first to report pre-gastrulation, circular expression of Brachyury. It will be interesting in the future to understand whether the epiblast border cells undergo a form of epithelial-to-mesenchymal transition to enable lumen closure. Thus, future studies of cell behaviour and transcriptomics will be required. Additionally, a comparison of signalling dynamics between species with and without epiblast lumenogenesis is needed to shed light into the role the lumen for subsequent development.

Epiblast lumenogenesis has not only evolved in chameleons, because *Mabuya mabouya*, a viviparous skink species, also exhibits also an epiblast lumen.^48^ This leads to us to hypothesise that epiblast lumenogenesis and pre-gastrulation amniogenesis may have evolved multiple times similar to viviparity^49^, especially considering the evolutional distance of 225MYA between veiled chameleon and *Mabuya mabouya*.^50^

## Supporting information

combined supplementary Figures

## Contributions

AW conceived of, planned, and carried out the project. NAS helped with experiments, data discussion, and project planning. HW carried out paraffin sectioning and Histology staining. MMC performed SEM and TEM. RK and AM carried out chameleon breeding and husbandry, ultrasounds together with AW and NAS and analysed the breeding date to lay date times. SAW provided critical feedback and mentoring. FH provided critical feedback and supervised AW. PAT supervised the project. AW wrote the manuscript with the help of NAS, SAW, FH, and PAT.

## Acknowledgements

The authors thank Dr Tom Kleist and Dr Cathy McKinney for their support with confocal imaging, and Dr Christopher White and Dr Sean McKinney for their help with confocal image processing. The authors thank all members of the Trainor and Williams labs as well as Dr Bonnie Kircher and Dr Thom Sanger for valuable discussions and feedback. We thank Dr Raul Diaz, Zoe Griffin, and Erika Pinto for cloning the *Bmp2*, *Wnt3a* and *Nr6a1 in-situ* probes. Research in the Trainor lab is supported by the Stowers Institute for Medical Research, and N.A.S is supported by a K99/R00 Pathway to Independence award from the National Institute of Child Health and Human Development (HD114881). AW is supported by a postdoctoral research fellowship of All Souls College, University of Oxford. AW and FH were supported by the Newton Trust, the European Research Council (695669) and the Engineering and Physical Sciences Research Council (EP/Y032756/1).

## Methods

### Data availability

Upon publication, original data underlying this manuscript can be accessed from the Stowers Original Data Repository at http://www.stowers.org/research/publications/xx.

### Limitations of working with a non-model organism

Many tools that are standard for model organisms such as mouse, chicken or Xenopus have not been established in chameleons. For example, no commercially available chameleon-specific antibodies exist. We tested a number of tissue marker antibodies, such as Oct4, Sox2, Cdx2, Gata6, and Gata4 that showed either no signal or expression patterns that diverge from what has been published for the species they have been raised against. Only for highly conserved genes such as Laminin and *β*-catenin could we detect signals that appears specific and were in accordance with protein localisation patterns in other species. Similarly, common tools to enable functional experiments such as transgenics or inhibitor treatments in *in vitro* cultured pre-oviposition embryos have not yet been established. As such, we cannot provide mechanistic proof for hypothesised cell and tissue movements.

### Chameleon Husbandry

Chameleons were housed in accordance with SIMR IACUC protocols 2022-147 Lizards and Snakes (Squamata) and 2023-160 Squamate Development following previously published husbandry protocols^20,51^. Stowers Institute for Medical Research is an AAALAC accredited institution. Partially screened enclosures measuring approximately 2’x2’x4’ are used for housing and are cleaned daily. Mercury vapor and T5 lighting provides access to UV sources as well as basking opportunities. Each enclosure contains a variety of artificial and natural vines, branches and foliage. Animals are fed once daily (a variety of insects and vegetables are provided on a rotating basis) while dietary supplementation is provided every other feeding. Misting and whole room humidification occurs several times daily. Adult chameleons are housed individually except during documented mating events. Breeding age females are provided lay tubs containing a sand/soil/peat moss mixture. These tubs are misted as needed to maintain appropriate moisture levels. Eggs are removed for artificial incubation within 24h of oviposition.

### Ultrasound

Ultrasound was performed as non-invasive method to track folliculogenesis and egg maturation using the Visual Sonics Vevo 1100 ultrasound system with a MS550D probe. For this, the chameleon was carefully manually restrained. Conductive gel was placed on the lower abdomen and then ultrasound images were taken. A minimum of 3 images were taken per side. Eggshells could be distinguished by their lighter colour as shown in Figure S2. This procedure was carried out every 48h to monitor eggshell maturation.

### Chameleon Euthanasia

Female chameleons were anesthetized using isoflurane until the toe pinch reflex was absent. For euthanasia 50% (v/v) of tricaine methane sulfonate (MS-222) solution was injected into the heart (1 ml per 250-300g body weight). After the loss of coloration and apparent death, we performed secondary euthanasia in the form of heart removal (pneuomothorax). This method of euthanasia is consistent with American Veterinary Association (AVMA) guidelines.

### Chameleon Dissection

The skin was opened along the midline and pinned to the sides. Then the abdominal cavity was opened and the two uterine horns containing eggs were isolated and each moved to PBS. Then, the ovaries were dissected and fixed in 4%PFA. In case no ovulation had taken place yet, the ovaries containing vitellogenic follicles were isolated and moved to PBS. All dissections were carried out in PBS.

### Embryo Dissection

In **cleavage-stage embryos**, the eggshell has not matured yet and the embryonic shield can be distinguished through the shell as white oval on the yolk. The eggshell is carefully cut in half and removed from the yolk which at this stage is still quite solid. The yolk was then turned until the embryonic shield was facing upwards. Then, the embryonic incisions were made around the embryonic shield and the shield overlaying a layer of yolk for stability as the embryo is not cohesive yet isolated through needles from the remaining yolk.

**Post cleavage but pre-lumenogenesis embryos** are embedded in the omphalopleure, which has grown around the yolk. The volume of the yolk is slightly lower than the volume of the eggshells at these stages, which enables removal of the eggshell halves without damaging the yolk. The embryo can be distinguished from the remaining yolk as beige-coloured circle or oval. The yolk is carefully positioned until the embryo is facing upwards, then the omphalopleure surrounding the embryo is cut using scissors and the embryo peeled off through forceps. The embryo is in direct contact with the yolk in contrast to chicken embryos. For **mature pre-oviposition embryos**, the omphalopleure has attached to the eggshells. This became apparent when removing the two eggshell halves from the yolk which had a rougher, less clean surface. The eggshells were held with one pair of forceps while a second pair of forceps was moved carefully along the eggshell to separate omphalopleure from the eggshell, once a portion of the omphalopleure separated, the membrane was pulled of the eggshells using forceps. The membrane was stretched out and the embryo located as solid oval or circle that was surrounded by a ring of denser cells. The embryo was cut out of the omphalopleure using needles and moved into fixative.

For **0dpo embryos**, the eggs were cleaned and then cut along the long axis into two halves. As the eggs are under pressure it is important to be careful when opening the shell. After the yolk was removed, the shells were placed into PBS. From here onwards, the procedure for mature pre-oviposition embryos was followed.

All embryos were moved into ice-cold 4% (v/v) PFA in DEPC treated PBS and fixated overnight at 4°C. A set of embryos from each clutch was retained in PFA and used for immunofluorescence stainings while the rest was washed 3x in PBS on the following day and dehydrated into MeOH (25/50/75/100/100%(v/v)) and stored at -20°C to preserve RNAs for subsequent analyses.

### DNA extraction of germinal vesicles

Germinal vesicles were dissected from the embryonic plates of oocytes and collected in one tube in PBS. A second tube was filled with some follicular tissue as control. Following dissection, the vesicle were spun down in a table-top centrifuge and the maximum amount of supernatant was removed. Then, 50ul of solution A to germinal vesicles/100ul to follicles (10N NaOH, 0.5M ETA in H2O) were added and incubated for 50min at 95°C. The tubes were cooled down and spun down and then 50ul (germinal vesicles)/10ul (follicles) of solution B was added and mixed. The DNA was stored at -20°C. The follicles were diluted 1:5 in 10mM Tris pH8, the germinal vesicles were kept as is.

### Genotyping

For genotyping two male loci we used previously published M2 and M3 primer pairs.^52^ 50% of fertilized zygotes would be male. No oocytes will have male-specific markers. We used the following two sets of primers for autosomally located *Hox* genes (*HoxB8* and *HoxB9*) as positive controls.

*HoxB8* F 5’-CGGCTTTGTACTTGGAGAAGA

*HoxB8* R 5’-GTGGAGTACGTGGAAACCAATA

*HoxB9* F 5’-GGGATACCCACCAAACTCTATC

*HoxB9* R 5’-AAATCCAGAGACCGCCATATC

The samples were subjected to PCR together with male and female control DNA as well as one empty control ((10ul 2x PhireMastermix, 1ul Primer F (10μm), 1ul Primer R (10μm), 1ul DNA, 7ul H2O) run as follows 5min 98°C, 35x(98°C 5sec, 64.8°C 5sec, 72°C 30sec), 72°C 1min. The PCR products were run on a 1% agarose gel in TAE buffer and developed on a chemismart.

### Immunofluorescence Staining

Embryos were placed on a rocker for all wash and incubation steps. Embryos were washed 3x in PBS and then permeabilised for 20min in permeabilisation buffer (0.3M TritonX100, 0.1M glycine in PBS), rinsed 2x in PBST (PBS, 0.1% Triton X100) and then incubated in blocking solution (0.1% Triton X-100, 1% v/v donkey serum in PBS) for 1h at RT. Primary antibody incubation was carried out overnight at 4°C. Next morning, the samples were washed 3x in PBST for 10min at RT and then incubated in secondary antibodies plus DAPI in blocking solution for 2-3h at RT or overnight at 4°C in the dark. The embryos were washed 3x in PBST in the dark and then equilibrated through a glycerol series (25%/50% glycerol in PBST) for 15min each at RT. The embryos were mounted in VECTASHIELD mounting medium (Vector Laboratories H-1200-10) between two cover slips to enable imaging from dorsal and ventral sides using vacuum grease using a 20x air objective on a Leica SP8 or a 20x air objective on a LSM800 or LSM 900 confocal microscope. For tile scans, 15% stitching overlap was used. The nucleoli of the germinal vesicles were imaged on a Nikon Eclipse Ti2 microscope equipped with a Yokagawa CSU W1 10,000 rpm Spinning Disk Confocal with 50μm pinholes using a 100x oil objective.

Primary antibodies: *β*-Catenin (rabbit, 1:200), Brachyury (goat, 1:200), Laminin (rabbit, 1:100). Secondary antibodies and fluorescent stainings: DAPI, donkey-anti-mouse-IgM-546, donkey-anti-mouse-IgG-488.

### Histology

Embryos that had been previously stored in 100% MeOH were stepwise rehydrated (100/70/50/30 5 min each at RT) then dehydrated to 70% EtOH (30/50/70 5 min each at RT). A couple drops of Eosin were added to the 70% to lightly dye the embryos to aid in orientation at embedding. Embryos were paraffin processed (Milestone, Pathos Delta Microwave Tissue Processor) with the following protocol w/o pressure: 70% EtOH 4 min at RT, 100% EtOH 5 min at 37C, Isopropyl Alcohol 5 min at 45C, Paraffin 15 min at 62C. After processing embryos were embedded in paraffin wax (Cancer Diagnostics, PureAffin® R56) and sectioned at a thickness of 5 µm on a microtome (Leica RM2255). H&E staining was performed using an automatic stainer (DP360, Dakewe (Shenzen) Medical Equipment Co.) with ST Infinity H&E Reagents (Leica Biosystems Cat. 3801698). Slides were mounted with Cytoseal 60 mounting media (VWR, 48212-187).

Embryos were imaged using Olympus Slide Scanner on a 20x objective.

### RNA *in situ* Hybridisation Probes

Probes for *Lefty*, *Nodal1*, *Nodal2*, and *Cerberus* have been published and validated previously.^17^

To clone RNA *in situ* probes for *Brachyury, Wnt3a* and *Nr6a1*, we collected embryonic RNA from the stages when the genes were likely to be expressed. We used SuperScript III First-Strand Synthesis System for RT-PCR (Invitrogen 18080-051) for cDNA synthesis, utilizing 100 ng of RNA per reaction. We caried out both Oligo (dT) and randomer hexamers reactions. The two reactions were mixed, and that cocktail was used to clone gene fragments for RNA *in situ* hybridisation probes.

*Bmp2* probe is 608bp, and was cloned into the pGEM-T Easy vector. The sequences flanking the probe are as follows: F 5’-TGCTGGACACCCGGCTGATG and R 5’-TCAGCCCTCCACCACCATTT. *Wnt3a* probe is 500bp and was cloned into the Zero Blunt TOPO vector (Invitrogen 450245) using the following primers: F 5’-GCCATTGGCCATCAATATTCC and R 5’-GTCCTTCCAGCTTCGTTGT. *Nr6a1* probe is 760bp and was cloned into the Zero Blunt TOPO vector (Invitrogen 450245) using the following primers: F 5’-GCTGATCGAAGATGGGTACAA and R 5’-AGAAACACACACGGGAGAAG.

### RNA *in situ* Hybridisation

A protocol optimised for chameleon RNA *in situ* hybridisation was followed.^17^ All incubation steps unless otherwise stated were carried out on a rocker. In short, Embryos were dehydrated into 100% MeOH (25/50/75/100/100% 5min each at RT) and stored overnight at -20°C. Following day, the embryos were re-hydrated (75/50/25% MeOH 5min each at RT) and washed 2x 5min in PBST/DEPC. Embryos were treated 7min with proteinase K (10ug/ml) without shaking and were fixed again in 4%(v/v)PFA/DEPC, 0.1% (v/v) glutaraldehyde for 20min. embryos were rinsed 1x 2min and washed 2×5min in PBST to be transferred to 50% PBST/DEPC 50% pre-hybridisation buffer (50% formamide, 2% SDS, 2% blocking reagent (all v/v), 5x SSC pH7.0, 250ug/ml tRNA, 100ug/ml Heparin) for 10min at 68°C. embryos were then incubated 1×10min and 1×1h in 100% pre-hybridisation buffer at 68°C. Then the probes were added at 1ug/ml in Pre-hybridisation buffer and the embryos were incubated overnight at 68°C. Next day, the probes were removed and the embryos rinsed (2min) in prewarmed solution X (50% (v/v) formamide, 1% (v/v)SDS, 2x SSC pH7.0) followed by 4 washes in solution X for 30min each at 68°C. Then, embryos were washed in solution X:MABT (1:1) for 10min at 68°C and then moved to RT for all following steps. Embryos were rinsed 3x in MABT (2min) and washed 2×30min). Embryos were washed 1×1h in MABT + 2% (w/v) blocking reagent and then washed 1×1h in 20% (v/v) lamb serum, 2% (w/v) blocking reagent in MABT. Then, anti-DIG was incubated in 20% (v/v) lamb serum, 2% (w/v) blocking reagent in MABT at 4°C overnight. Next day, embryos were washed 3×5min in MABT and then washed 7×45min in MABT at RT. Embryos were moved to NTMT (100mM NaCl, 100mM Tris pH9,5, 50mM MgCl2, 1% (v/v)Tween2-, 2mM levasimole) and washed 4×10min. Embryos were then developed in the dark at RT in NTMT + 1;400 BCIP & NBT each). The developing reaction was stopped through PBST. Embryos were then moved to PFA and stored at 4°C. For each timepoint, a sense control was run in parallel to the antisense probes.

Probes for *Lefty*, *Bmp2*, *Nodal1*, *Nodal2*, *Brachyury*, *Cerberus* have been published and validated previously.^17^ The probes for *Nr6a1* and *Wnt3a* were prepared new for this manuscript.

### Scanning Electron Microscopy

Samples for scanning electron microscopy were prepared as previously published.^30^ Briefly, after fixation samples were processed using an TOTO method (tannic acid, osmium tetroxide, thiocarbohydrazide, and osmium tetroxide), dehydrated in a graded series of ethanol, and dried in a Tousimis Samdri 795 critical point dryer. After mounting on stubs, samples were imaged in a Zeiss Merlin SEM at 8kV using the SE2 detector.

### Transmission Electron Microscopy

Samples for TEM imaging were also prepared as previously published^30^, with secondary fixation with osmium tetroxide and *en bloc* staining with uranyl acetate before dehydration in a graded series of ethanol, propylene oxide as a transitional solvent, and infiltration with Hard Plus resin (Electron Microscopy Sciences). After embedding samples were cured at 60C. samples in blocks were imaged in a Bruker SkyScan 1272 microCT to identify regions of interest, sectioned at 80nm with a Diatome diamond knife and imaged in a Zeiss Merlin SEM at 26kV and 700pA with aSTEM detector.

### Brightfield Imaging

Embryos were placed in a petri-dish coated with Agarose and filled with PBS. Imaging was carried out on a Leica DFC550 microscope. Manual stacks were taken that were rendered using the Helicon software (Focus 5.3 software).

### Image Processing

#### SP8

The Imaging shown in Figure 1F, S1B was performed using bidirectional scanning, which resulted in in every odd scan-line being shifted by 4 pixels. This happened due to a misalignment of the two scanners. The images were then corrected computationally by shifting every odd scan-line by 4 pixels to align with the even lines. The simple correction were applied in python.

### Image Analysis

All image analysis was carried out in Fiji, all graphs were prepared through GraphpadPrism10. For all measurements, the ROIs were saved to enable traceability of the measurements.

## Supplementary Information

**Figure S1: Oocyte morphology of the veiled chameleon.**

**A.** Brightfield imaging. Left: ovary filled with vitellogenic follicles at 30dpc. The uterine horn is visible beneath the ovary (arrows). middle/right: mature vitellogenic follicles inside the ovary with germinal discs (square) and germinal vesicles visible (asterisks) at 30dpc(middle) and 32dpc (right). Scale bars left 2cm, middle/right 2mm. **B.** Confocal imaging of DAPI stained germinal discs. Z-stack through the germinal vesicle, z10/40/66 are shown. The middle of the germinal vesicle is DAPI positive, arrows highlight DAPI-positive region. Scale bar 100μm. **C.** maximum intensity projection of spinning disc imaging of DAPI-positive centre of 2 germinal vesicles. DNA has segregated into distinct foci. Scale bars 10μm. **D.** Chameleon handling for ultrasound of abdominal cavity.

**Figure S2: Following folliculogenesis and eggshell maturation via ultrasound**

A-F. Ultrasound imaging of veiled chameleon abdomen. A. immature vitellogenic follicles arrows indicate follicles. B. mature vitellogenic follicles. Arrows indicate border between follicles which is dark indicating no shell development yet. C. initiation of eggshell formation. arrows indicate the thin eggshells in-between eggs. D. early eggshell. Arrows indicate eggshell, asterisks point out space between yolk and eggshell characteristic for early shelling stages. E. maturing eggshell. Arrows indicate thickened eggshell, asterisks the space between yolk and shell. F. mature eggshell on day before oviposition. Arrows indicate eggshell, no empty space between yolk and eggshells at mature stages. asterisks indicate uterine wall. All scale bars 1mm.

**Figure S3: Initial cleavage division pattern**

A. Ultrasound imaging of eggshells of the eggs dissected for Figure 2B-C, SB/C. arrows indicate the very thin eggshells. scale bars 2mm. B. SEM imaging of initial embryo cleavage. blue highlight indicates area of embryonic plate. Squares indicates regions of zoom-ins. (i) cleavage furrow. Zoom-in on initial cleavage (i*) cleavage initiation. (ii/ii*) embryo border. Blue indicates embryonic plate, arrows point out embryo border. (ii**/***) furrow initiation. arrow indicated hole that facilitates cleavage initiation at the embryo border. Scale bars: initial cleavage=400μm, i/ii=100μm, i*/ii*=20μm, ii**=40μm, ii***=5um. **C.** TEM of initial cleavage stage blastomere. Squares indicated regions of zoom-ins. (i) blastomere localisation. (i*) zoom-in on nucleolus. (i**) nuclear pores. Arrows indicates nuclear pores. (ii) mitochondria. Asterisks indicate regions of mitochondria. Squares indicate regions of zoom-in. (ii*) golgi apparatus. Arrows indicate golgi. (ii**/iii-iv) filaments. Arrows indicate differently-angled filaments in cytoplasm. scale bars: initial blastomere=100μm, i=40μm, i*/iv=100nm, i**=300nm, ii=3um, ii*/**/iii=200nm. **D.** SEM of blastomere formation. Squares indicate regions of zoom-ins (i) early blastomeres in middle of embryonic plate. (i*) cellularisation. Blue highlight mature cleavage furrow. (i**/***) cleavage furrow. Furrow highlighted in blue. (ii) furrow initiation. arrows indicate initiating furrow. scale bars: blastomere formation=200μm, i=100μm, i*=20μm, i**/ii=10μm, i***/ii*=2um. **E.** Ultrasound imaging of eggshells of eggs dissected for Figure 2D/S3D). arrows indicate eggshell. all scale bars=2mm.

**Figure S4: Quantitative analysis of cell shapes during late cleavage stages.**

**A.** Analysis surface area of embryonic cells of areas 1-4. Analysis of 3 embryos per area. Combined plot shown in Figure 2H. Scatter plot plus mean±SEM. Statistical Analysis One-way Anovas for each embryo p-value<0.0001. unpaired t-tests for each embryo area 1-2/2-3/3-4 p-value<0.0001. statistical analysis for combined data in main figure 2H. **B.** Analysis surface perimeter of embryonic cells of areas 1-4. Analysis of 3 embryos per area. Combined data on the right. Scatter plot plus mean±SEM. Statistical Analysis One-way Anovas for each embryo p-value<0.0001. Unpaired t-tests for each embryo and combined embryos area 1-2/2-3/3-4 p-value<0.0001 except for embryo 3 area2-3, here p-value 0.0039. **C.** Analysis cell circularity of embryonic cells of areas 1-4. Analysis of 3 embryos per area. Combined plot shown in Figure 2I. Scatter plot plus mean±SEM. Statistical Analysis One-way Anovas for each embryo p-value<0.0001. unpaired t-tests for each embryo area 1-2/2-3/3-4. P-values: Embryo 1 (1-2) =0.0003, (2-3)=0.4937, (3-4)=0.0563. Embryo 2 (1-2)<0.0001, (2-3)=0.1647, (3-4)=0.0434. Embryo 3 (1-2)<0.0001, (2-3)=0.1985, (3-4)=0.1542. statistical analysis for combined data in main figure 2I. **D.** Analysis three-cell vertices of areas 1-4. Analysis of 3 embryos per area. Combined plot shown in Figure 2K. Scatter plot plus mean±SEM. Statistical Analysis One-way Anovas for embryo 1/2 p-value<0.0001, embryo 3 p-value=0.0007. Unpaired t-tests for each embryo area 1-2/2-3/3-4. p-values: Embryo 1 (1-2)=0.4308, (2-3)=0.0057, (3-4)=0.5422. Embryo 2 (1-2)=0.2555, (2-3)=0.0039, (3-4)=0.6229. Embryo 3 (1-2)=0.4645, (2-3)=0.4330, (3-4)=0.0211.

**Figure S5: The stages of chameleon pre-oviposition development**

Table with images or schematics depicting each stage of development accompanied by short descriptions

**Figure S6: Analysis of cell shapes in embryos following cell-cell adhesion**

**A.** H&E staining of cross sections of 3 embryos following cell-cell adhesion. squares indicate regions of zoom-in. asterisks indicate holes in the hypoblast layer. arrows indicate epiblast layer. All scale bars:200μm. **B/C.** Quantitative analysis of dorsal to ventral cell populations of embryos shown in (A). (B) Cell area. Scatter plot plus mean±SEM. Statistical Analysis unpaired t-tests dorsal-ventral p-values (1)=0.0007, (2)=0.1887, (3)=0.0500. (C) Cell circularity. Scatter plot plus mean±SEM. Statistical Analysis unpaired t-tests dorsal-ventral p-values (1)=0.0016, (2)=0.4391, (3)<0.0001. D. Immunofluorescence staining of β-Catenin in embryo post cell-cell adhesion, maximum intensity projection. Squares indicate areas of zoom-in. Scale bar=200μm.

**Figure S7: Analysis of Cell shapes during Lumenogenesis**

**A-C.** Quantitative Analysis of Cell Circularity during lumenogenesis (embryos i-vi) in (A) central cells, (B) cells in ring surrounding epiblast centre, (C) outer cells. Scatter plot plus mean±SEM. Statistical Analysis One-way Anovas, all p-values<0.0001. **D-I.** Quantitative analysis of cell area in middle (inner) versus peripheral (outer) cells during lumenogenesis stages i-vi (corresponding to D-I). Scatter plot plus mean±SEM. Statistical Analysis One-way Anovas, all p-values<0.0001.

**Figure S8: Expression patterns of anterior-posterior patterning and gastrulation regulators.**

**A-E.** whole mount in situ hybridisations at subsequent stages of development from stage VIII-XIX of (**A**) *Cerberu*s, (**B**) *Lefty*, (**C**) *Nodal1*, (**D**) *Bmp2*, (**E**) *Brachyury*. Stages are annotated on top of each column. All scale bars 200μm.

**Figure S9: Brachyury protein expression patterns**

**A.** Maximum intensity projection of F-actin (top row) and Brachyury (bottom row). Embryo stages defined on top of each column. All scale bars 100μm.

## Notes

### Competing Interest Statement

The authors have declared no competing interest.

