## Supplementary figures and images for "Epiblast Lumenogenesis is not a mammalian-specific trait"

### combined supplementary Figures

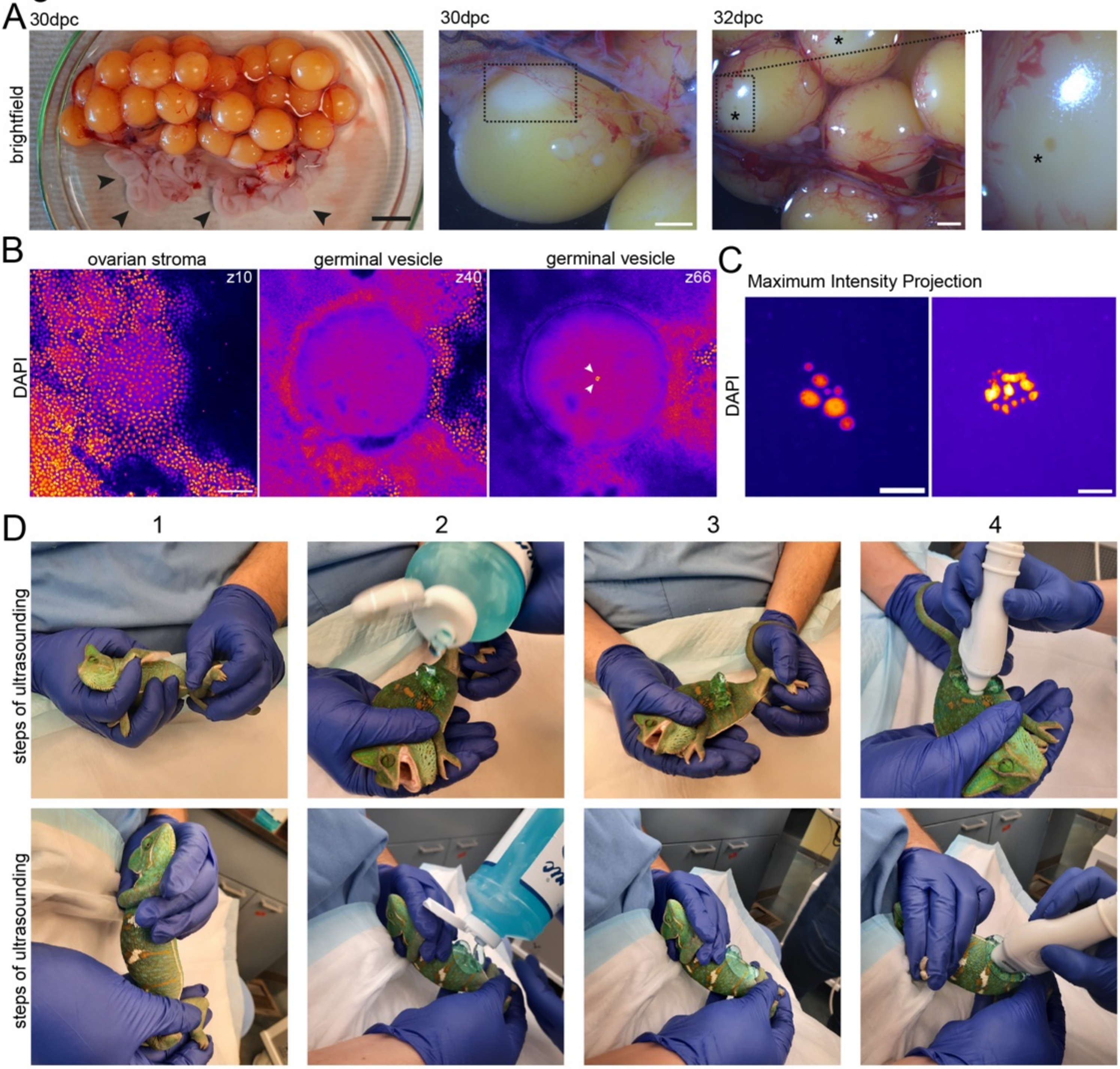

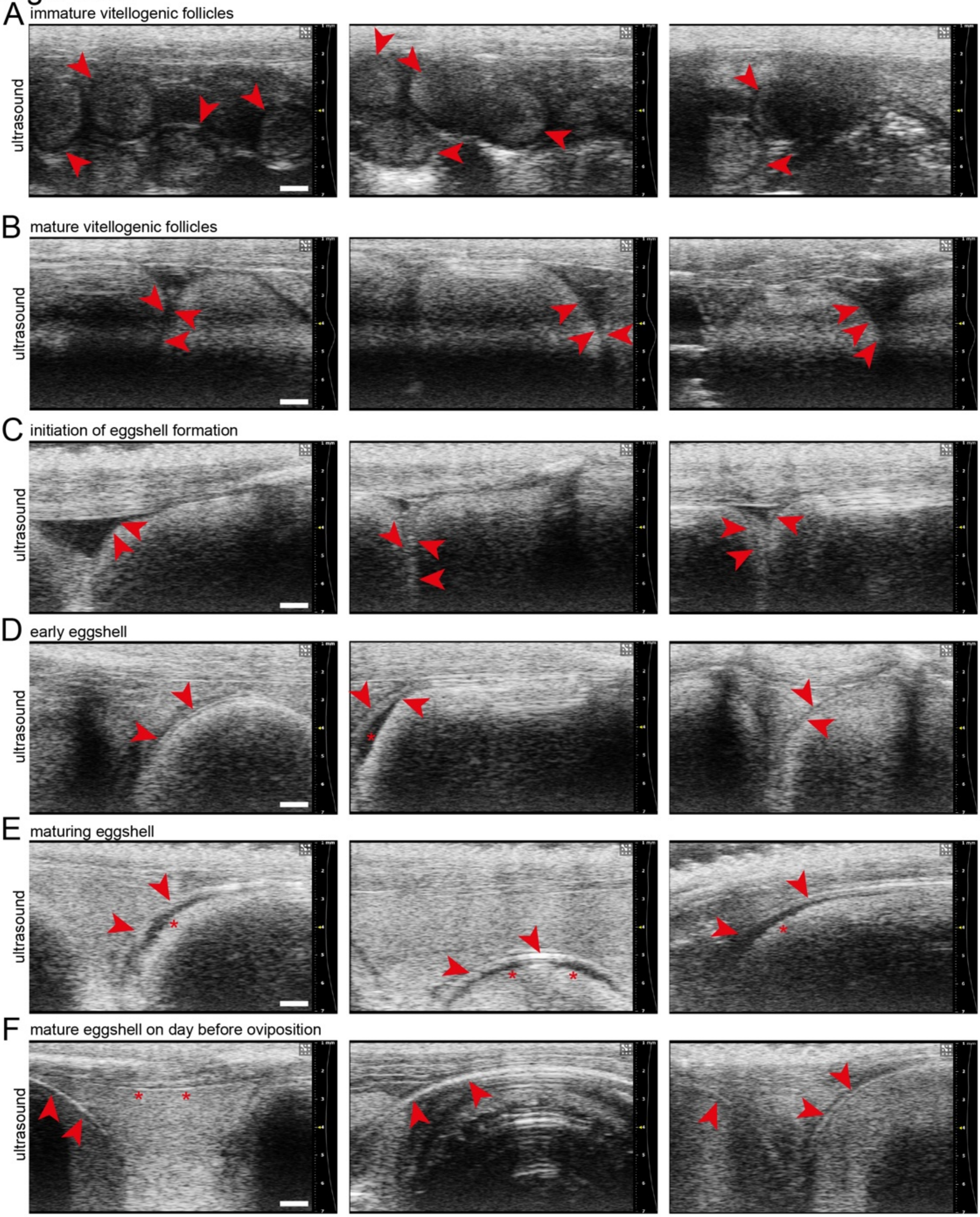

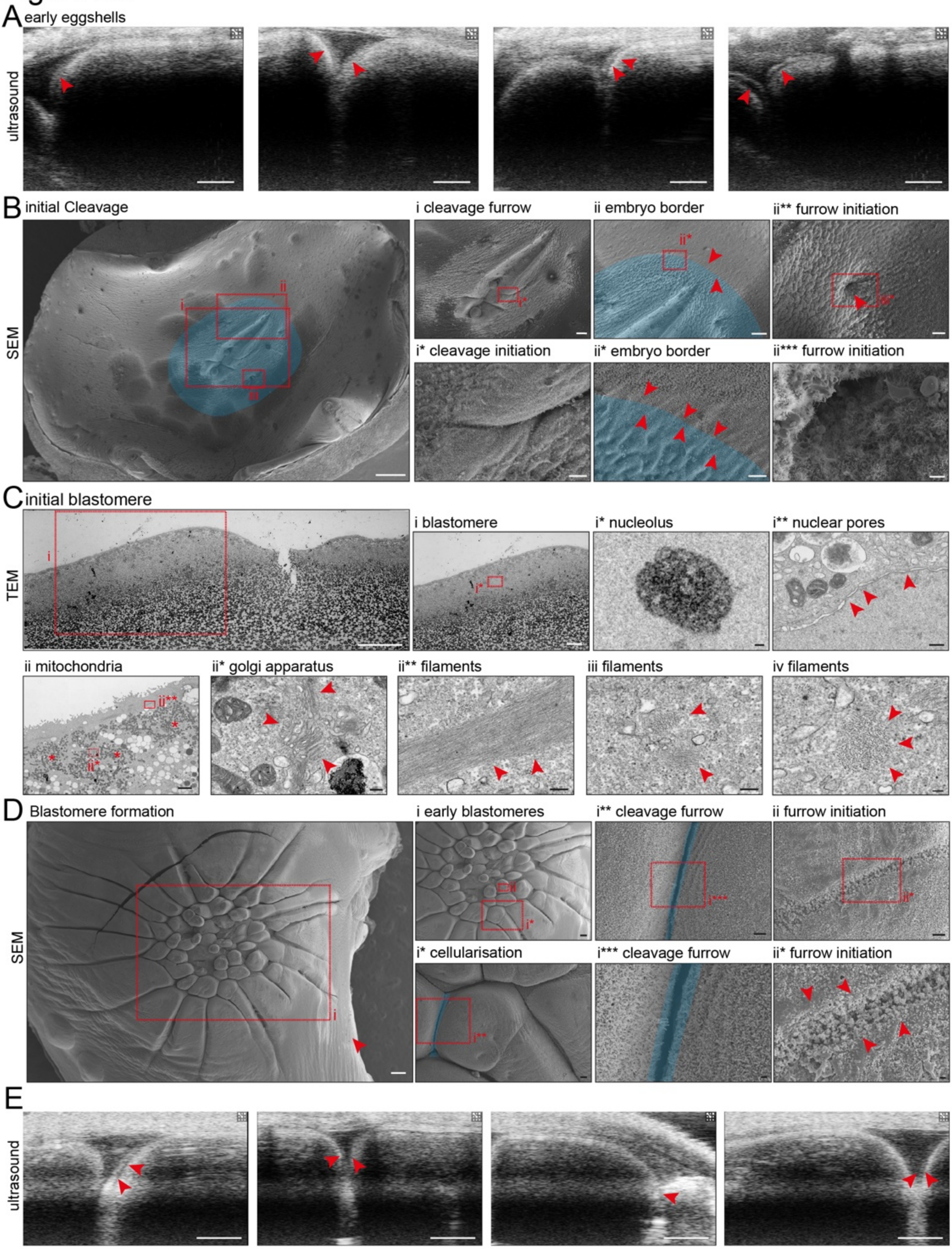

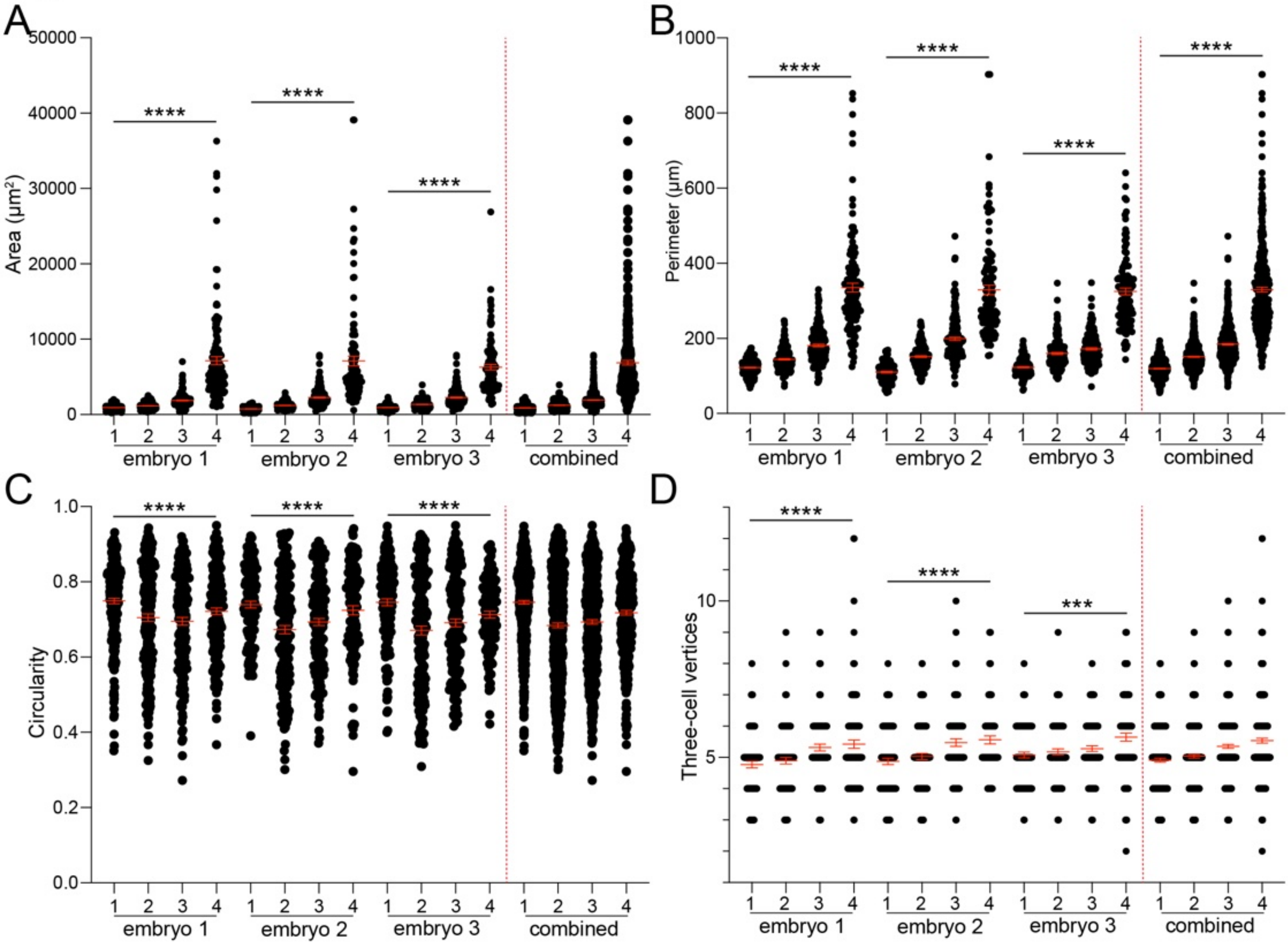

#### Figure S5 - stages of embryogenesis

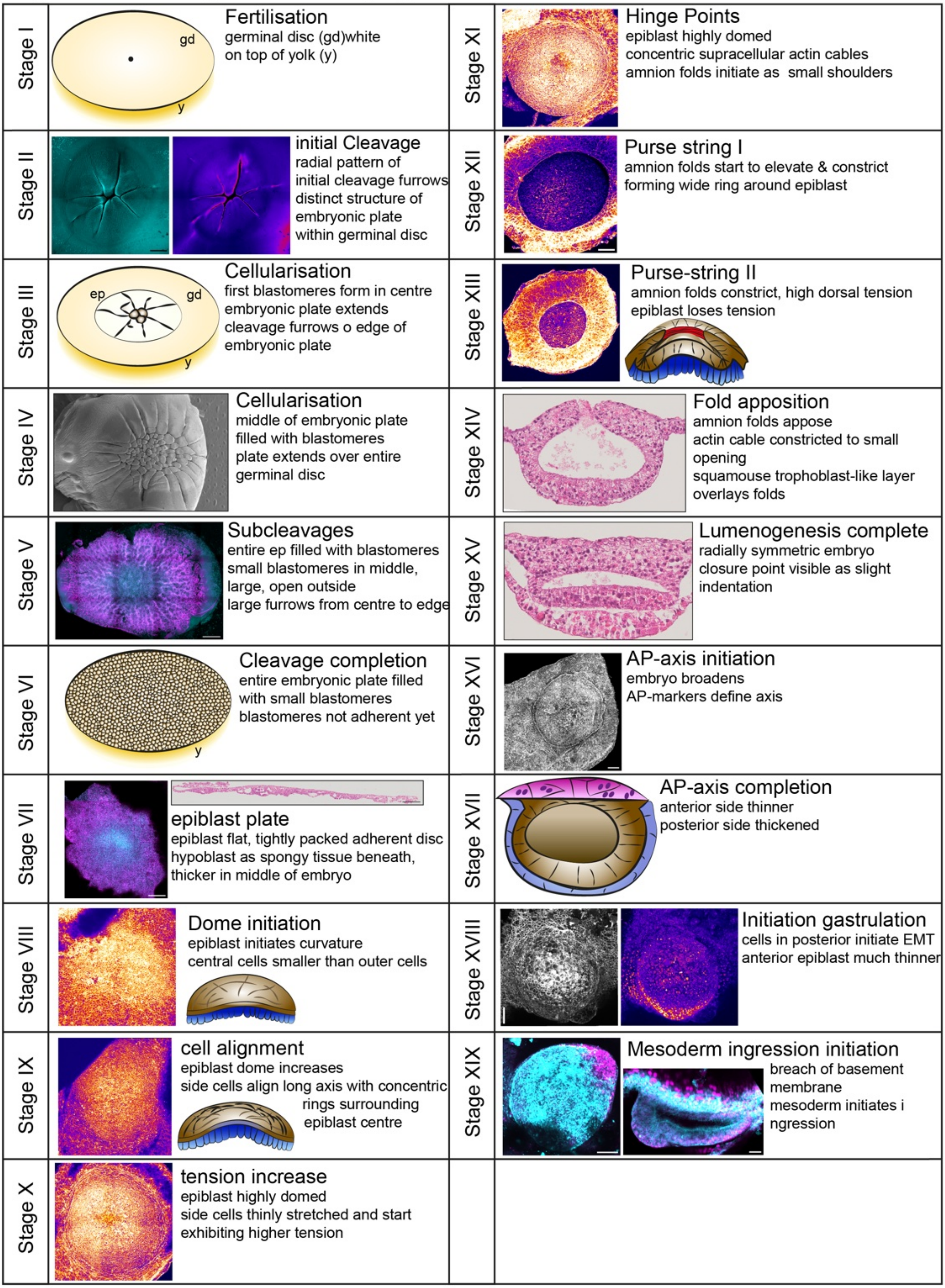

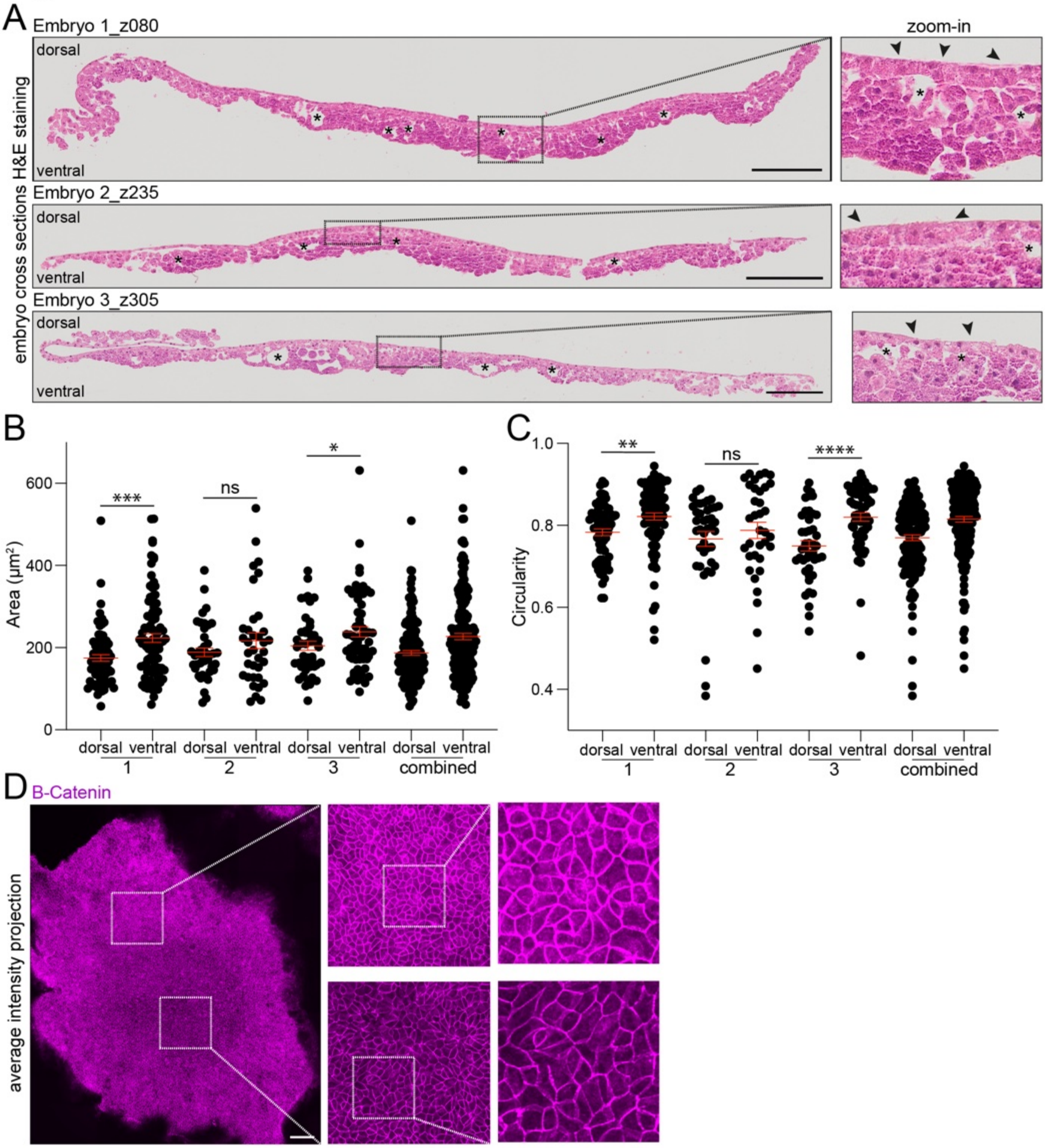

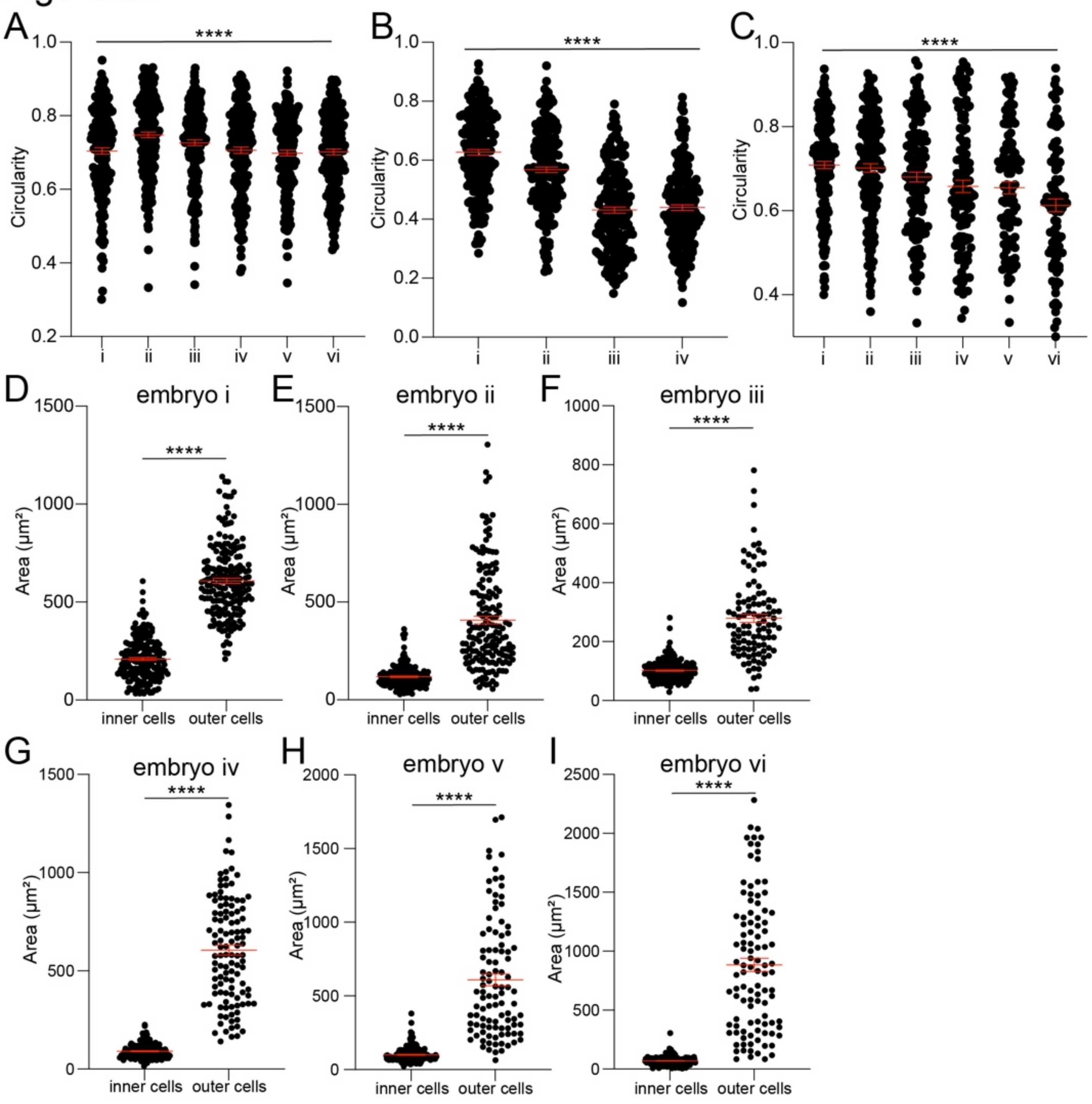

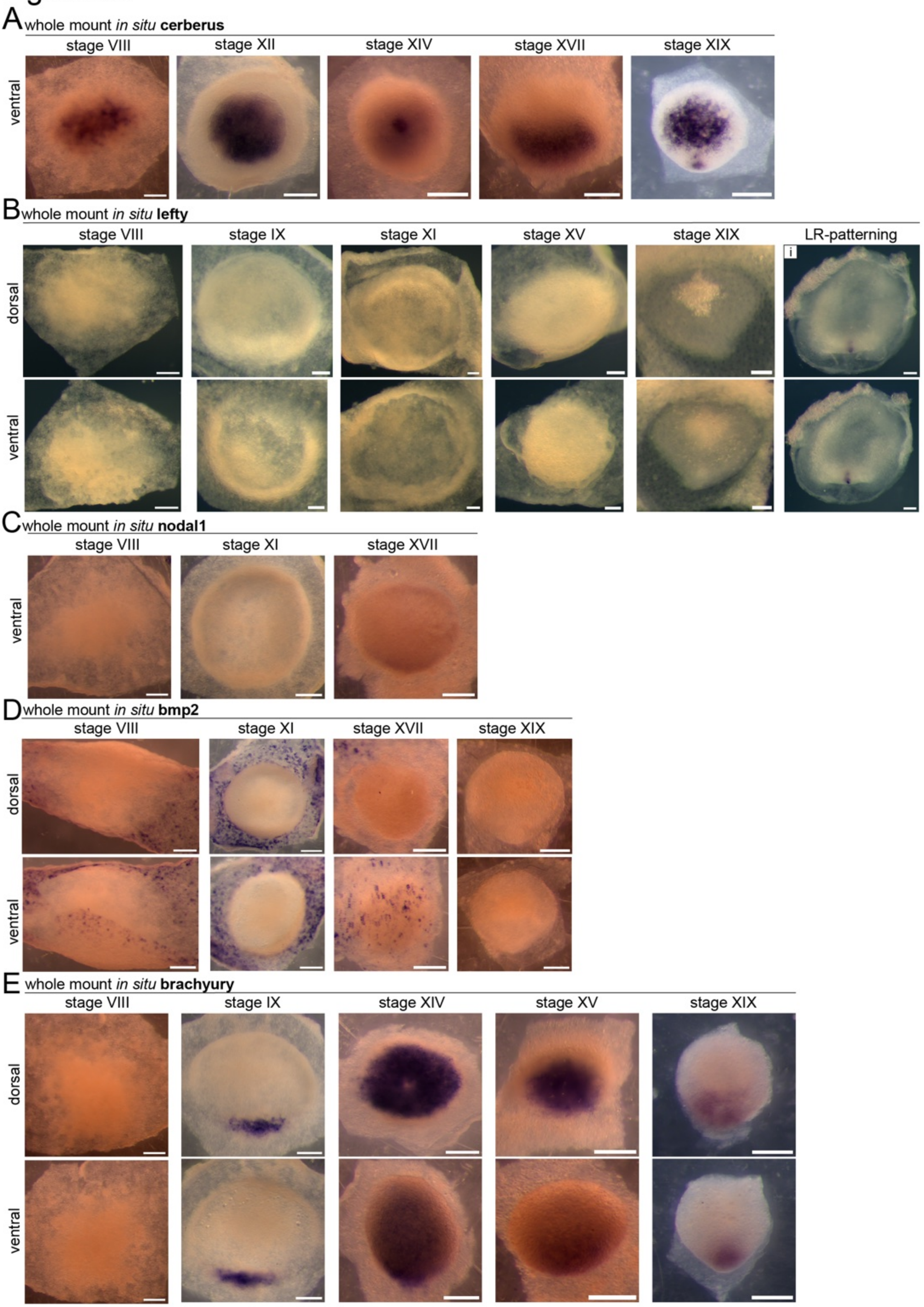

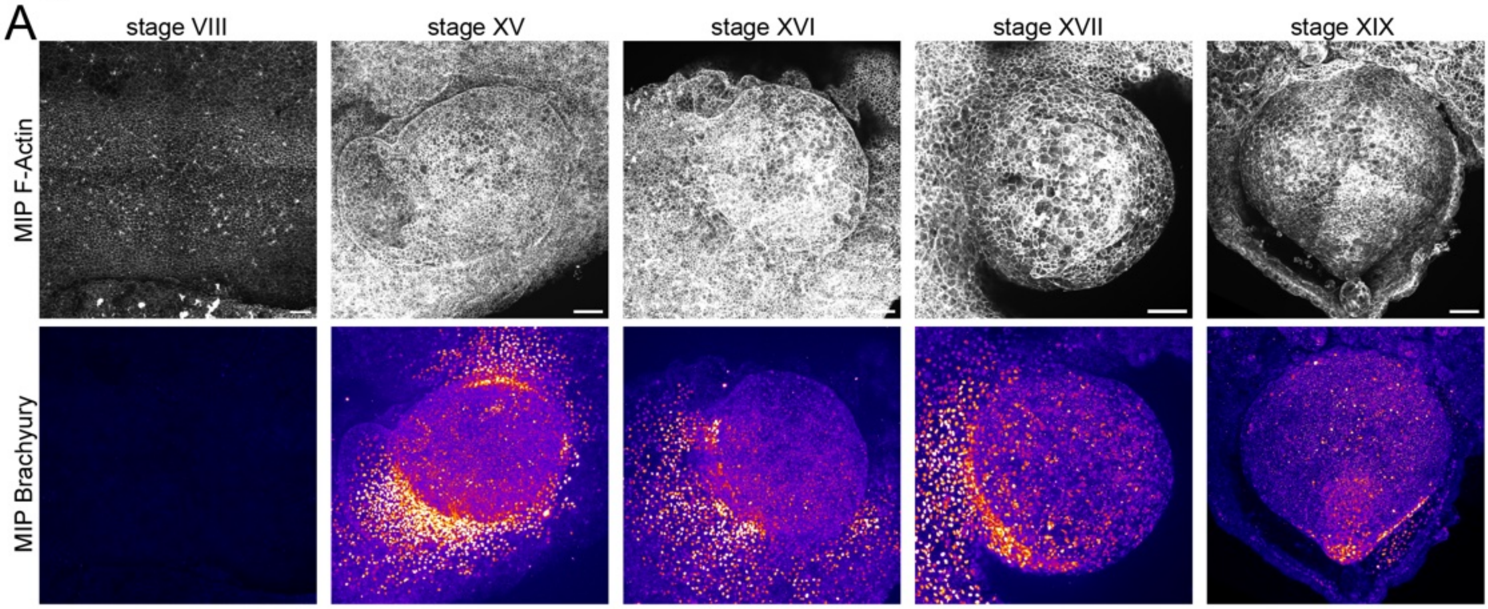
